# Leveraging phenotypic variability to identify genetic interactions in human phenotypes

**DOI:** 10.1101/2020.07.28.225730

**Authors:** Andrew R. Marderstein, Emily Davenport, Scott Kulm, Cristopher V. Van Hout, Olivier Elemento, Andrew G. Clark

## Abstract

While thousands of loci have been associated with human phenotypes, the role of gene-environment (GxE) interactions in determining individual risk of human diseases remains unclear. This is partly due to the severe erosion of statistical power resulting from the massive number of statistical tests required to detect such interactions. Here, we focus on improving the power of GxE tests by developing a statistical framework for assessing quantitative trait loci (QTLs) associated with the trait means and/or trait variances. When applying this framework to body mass index (BMI), we find that GxE discovery and replication rates are significantly higher when prioritizing genetic variants associated with the variance of the phenotype (vQTLs) compared to assessing all genetic variants. Moreover, we find that vQTLs are enriched for associations with other non-BMI phenotypes having strong environmental influences, such as diabetes or ulcerative colitis. We show that GxE effects first identified in quantitative traits such as BMI can be used for GxE discovery in disease phenotypes such as diabetes. A clear conclusion is that strong GxE interactions mediate the genetic contribution to body weight and diabetes risk.

## Introduction

In Cappadocia, Turkey, traces of an asbestos-like, cancer-causing fiber was found in the materials of villagers’ homes and was prevalent in the air. However, this alone could not explain an epidemic where 50% of all Cappadocia villagers died from mesothelioma, compared to only 4.6% of asbestos miners with at least 10 consecutive years of work^1^. After nearly three years of living amongst the villagers, Roushdy-Hammady *et al*. documented a Cappadocia villager pedigree and described a highly penetrant Mendelian transmission of disease^2^. Once the pathogenic *BAP1* mutations were found^3^, follow-up experimental studies^4-6^ illuminated how *BAP1* and asbestos exposure synergistically cause dangerous oncogenic effects in a gene-environment interaction (GxE)^7^.

This example is one of only a few well-characterized GxE interactions in humans, which have mostly appeared as modulators of Mendelian disorders’ penetrance. In human genetics, the primary focus has been characterizing the average relationship between individual genetic variants and a phenotype. While we have identified thousands of associations across a spectrum of human phenotypes at the single variant level^8^, research in model organisms and cell cultures have constantly shown that genetic effects are context-dependent^9-13^. As one example, the genetic effects governing lifespan in *Drosophila melanogaster* within one environment do not alter lifespan within another environment^10^. The incomplete penetrance for many common human diseases, such as the *APOE* E4 allele on Alzheimer’s disease risk or smoking on lung cancer risk, imply important genetic and environmental modifiers of disease onset^14-17^.

While GxE interactions are expected to be numerous, it is debated how important these interactions are to human genetics^18-21^. If they play a significant role, identifying these interactions can enable more accurate genetic prediction^12^, especially at the individual level. Current state-of-the-art prediction models use polygenic scores (PGS)^22^, which combine additive effects of genetic variants into a single risk measure. Clinical and environmental factors are used to improve model prediction, but potential interactions with genetic information are not commonly considered. Additionally, PGS have poor transferability across populations^23^, possibly driven by environmental factors. If genetic effects on human phenotypes vary from person-to-person due to interactions, then more individualized prediction could be realized by first identifying genetic interactions and estimating their effects.

Because there is modest power to detect interactions in large human population cohorts, efficiently identifying the interactions remains an important statistical and computational challenge. To address these difficulties, we make use of a previously characterized observation that most GxE interactions with large effect size can be revealed as a change in the variance of a quantitative phenotype during a one-SNP-at-a-time genome-wide association study (GWAS) (**Fig 1**)^24-26^. This insight lets us identify strong GxE interactions associated with a given quantitative trait using a two-step approach. First, we look for genome-wide SNPs that are associated with the variance of the trait, thus identifying what are known as variance Quantitative Trait Loci (vQTLs). Second, we use these vQTLs to screen for potentially strong GxE interactions associated with the same phenotype. Scanning for vQTLs involves just a single test per SNP, so it provides a powerful inroad for discovering genetic interactions by nominating loci as promising candidates for an interaction.

**Figure 1:**
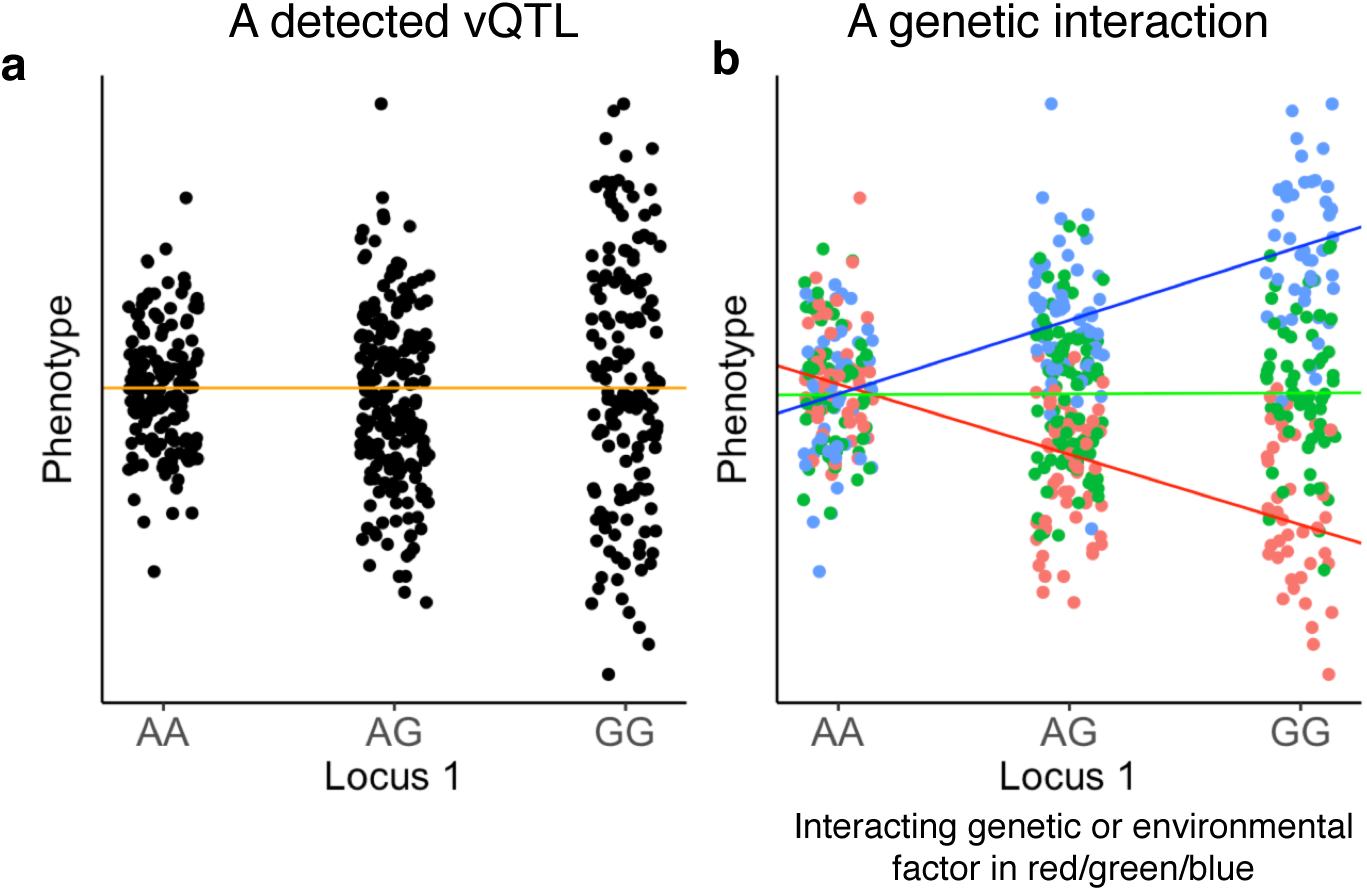
vQTLs could arise from a genetic interaction. (a) We refer to a genetic variant associated with the variance of the phenotype as a variance QTL (vQTL). The orange line, representing the line of best fit, has slope ≈ 0 and indicates that the mean of the phenotype does not change with a difference in genotype. (b) A vQTL could also arise from a genetic interaction. The displayed data in (b) is the same data as in (a), except the points are colored to reflect the genotype at a second locus or the level of an environmental variable. This second factor interacts with Locus 1 to create a mean-based interaction effect, and this mean-based interaction effect gives the appearance of a variance QTL at Locus 1. Data in both figures are simulated.

In the present study, we introduce a statistical framework to nominate SNPs for GxE interaction testing by leveraging differences in the means (muQTLs) and the variances (vQTLs) of a phenotype. We apply these methods to study the genetic basis of variation in body mass index levels (BMI)^27,28^. We further explore the role for interactions across human disease and perform in silico functional analyses for implicating relevant cell types and pathways, providing new insights into the architecture of human phenotypes.

## Results

### Deviation Regression Model discovers vQTLs that are due to GxE interactions

To identify genetic variants associated with the variance of quantitative phenotypes, we considered several tests (see Methods)^29-33^, including a new approach that we refer to as the Deviation Regression Model (DRM). In the DRM, a linear regression is performed on a single SNP and phenotype, where the minor allele count is used as the independent variable and the absolute difference between an individual’s phenotypic value and the phenotype medians within each genotype is used as the dependent variable (after covariate adjustment) (**Fig 2a**) (Methods). The effect sizes and *P*-values are used to estimate the variance effect of a SNP and assess vQTL significance, similar to a standard GWAS.

**Figure 2:**
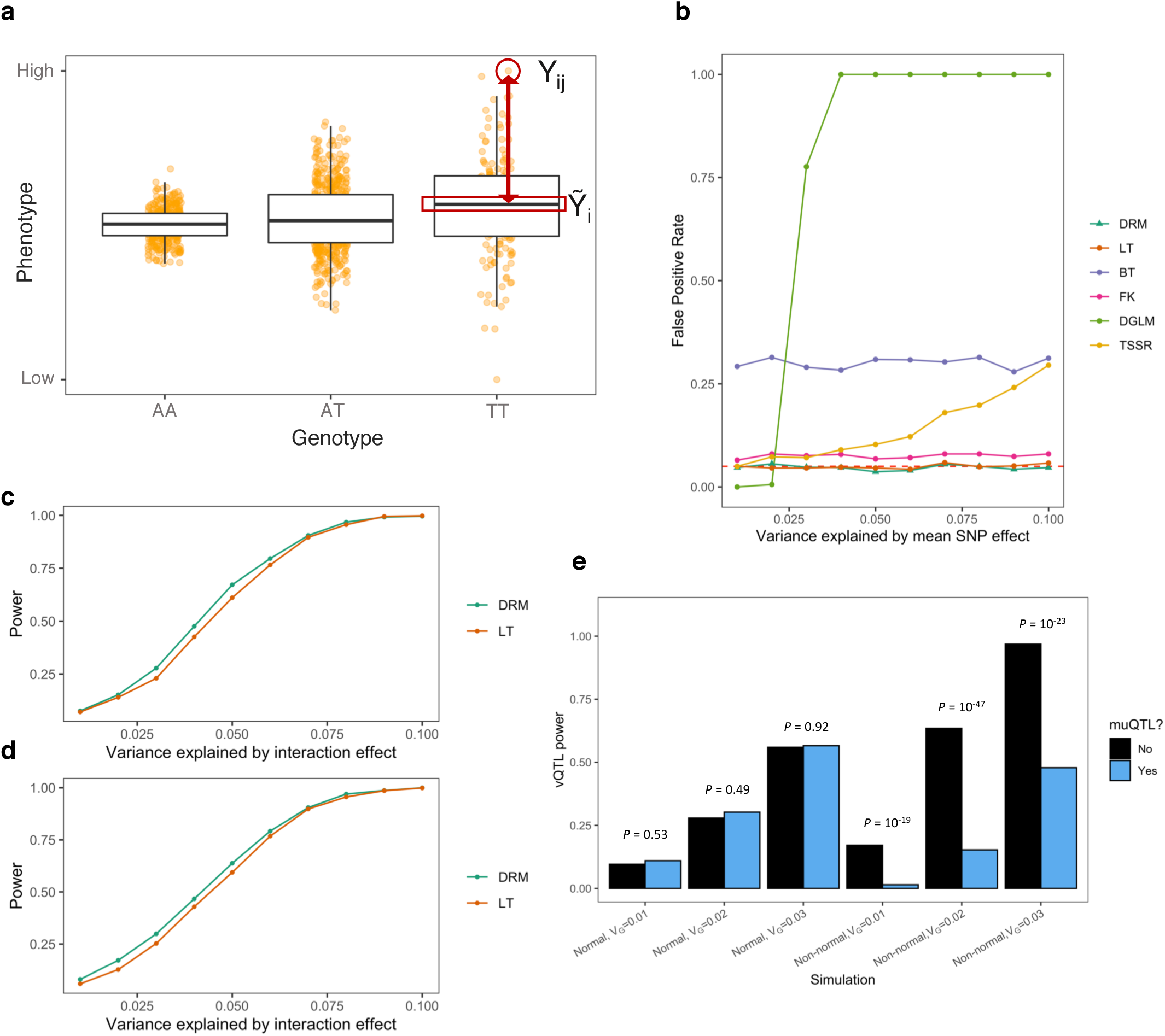
Assessing a variance test for finding SNPs with interaction effects. (a) The Deviation Regression Model uses the absolute difference between an individual’s phenotype Y_ij_ (for each genotype *i* and individual *j*) (*y*-axis) and the within-genotype phenotype median 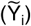 as a dependent variable. The absolute difference, Z_ij_, is modeled in a linear regression across genotypes (*x*-axis). Simulated data shown. (b) False positive rates for different variance tests at SNPs with varied mean effects in a non-normal phenotype: Deviation Regression Model (DRM), Levene’s test (LT), Bartlett’s test (BT), Fligner Killeen test (FK), double generalized linear models (DGLM), and a two-step squared residual approach (TSSR). (c-d) Power of the DRM and LT in normally distributed phenotypes (c) and chi-squared phenotypes (d). (e) vQTL test power, quantified by the DRM, stratified by whether the SNPs are detected by a muQTL test (linear regression). By using a 2-by-2 contingency table representing the counts of muQTL and vQTL test rejection across 1000 simulations, Fisher’s exact test assessed whether muQTL power and vQTL power show non-random association. *P*-values displayed.

We first used simulation to quantify the false positive rate (FPR) for the different variance tests. We tested the FPR using a scenario where a single SNP affects the mean of the phenotype, and thus a variance effect should not be detected except by random chance. We generated a SNP genotype and a phenotype value for 10,000 individuals. Across 1,000 simulations, we tested for a variance effect, and calculated the FPR as the proportion of simulations where the nominal *P* < 0.05. We found that the DRM maintained FPR near the expected 5% across trait distributions, demonstrating robustness of the DRM test to a SNP with only a mean effect on the phenotype. Levene’s test^33^ (LT) had similar performance. However, our simulations found extensive false positive rates for several other variance tests^29-32^, including the Fligner-Killeen test or the double generalized linear model (**Fig 2b**; Supp Fig 1a).

We further used simulations to test the ability of vQTLs to reveal SNP-by-factor pairs with an interaction effect on the phenotype. We repeated the previous FPR simulations, except generated an environmental factor that interacts with the SNP to influence the mean of the phenotype. As the interaction became stronger (variance explained by the interaction, V_G_, increases from 1% to 10%), the DRM’s power across 1000 simulations to detect an interaction effect at a SNP increased from less than 10% to 100% (**Fig 2c-d**). Additionally, we found an 11.9% power increase for the DRM compared to LT at smaller interaction effects (V_G_ ≤ 5%) (**Fig 2c-d**). Therefore, in our simulated data, vQTLs could identify SNPs with large-effect interactions and the DRM had superior power and improved FPR compared to other variance tests (Supp Note 1; Supp Fig 1-2).

Finally, we contrasted the use of vQTLs for identifying SNPs involved in an interaction with the use of mean-associated loci (muQTLs, as first identified using linear regression). Using a similar testing procedure, we simulated SNP-by-factor interactions where the direction of the SNP effect changes depending on the interacting factor (Methods). Across 1,000 simulations, we found that muQTL test power and vQTL test power were not positively correlated. For example, a vQTL test’s power at non-muQTLs was 62.9% when V_G_ = 2%, compared to 14.7% at muQTLs (**Fig 2e**). Our results show that positive and negative effects from a single SNP due to an interaction can remove a muQTL signal, and thus vQTL methods provide a complementary and robust approach to discover interactions (such as shown in **Fig 1**). Finally, although the muQTL approach had increased power to detect the causal SNPs compared to the vQTL approach (Supp Fig 1d), we note that muQTL approaches will pick up variants that directly impact the trait, but which are not involved in an interaction (leading to a high FPR). Therefore, our vQTL approach can identify SNPs involved in GxE interactions with higher specificity than using muQTLs.

### Genome-wide association studies in UK Biobank

Hundreds of variants have been associated with body mass index (BMI), highlighting that diverse pathways regulate body weight, from immune system activation to leptin signaling to the central nervous system^27,28^. Furthermore, environmental influences and lifestyle choices such as diet, exercise, and gut microbiome composition^34^ also have a major influence on BMI. Therefore, we hypothesized that there may be strong GxE interactions that regulate BMI, and these interactions may appear as a change in the BMI variance at a SNP. In 275,361 unrelated British European individuals from UK Biobank (UKB), we searched for genetic variants associated with the means (muQTLs) and variances (vQTLs) in untransformed body mass index (BMI) values (*P* < 5 × 10^−8^) (**Fig 3a**; Supp Note 2).

**Figure 3:**
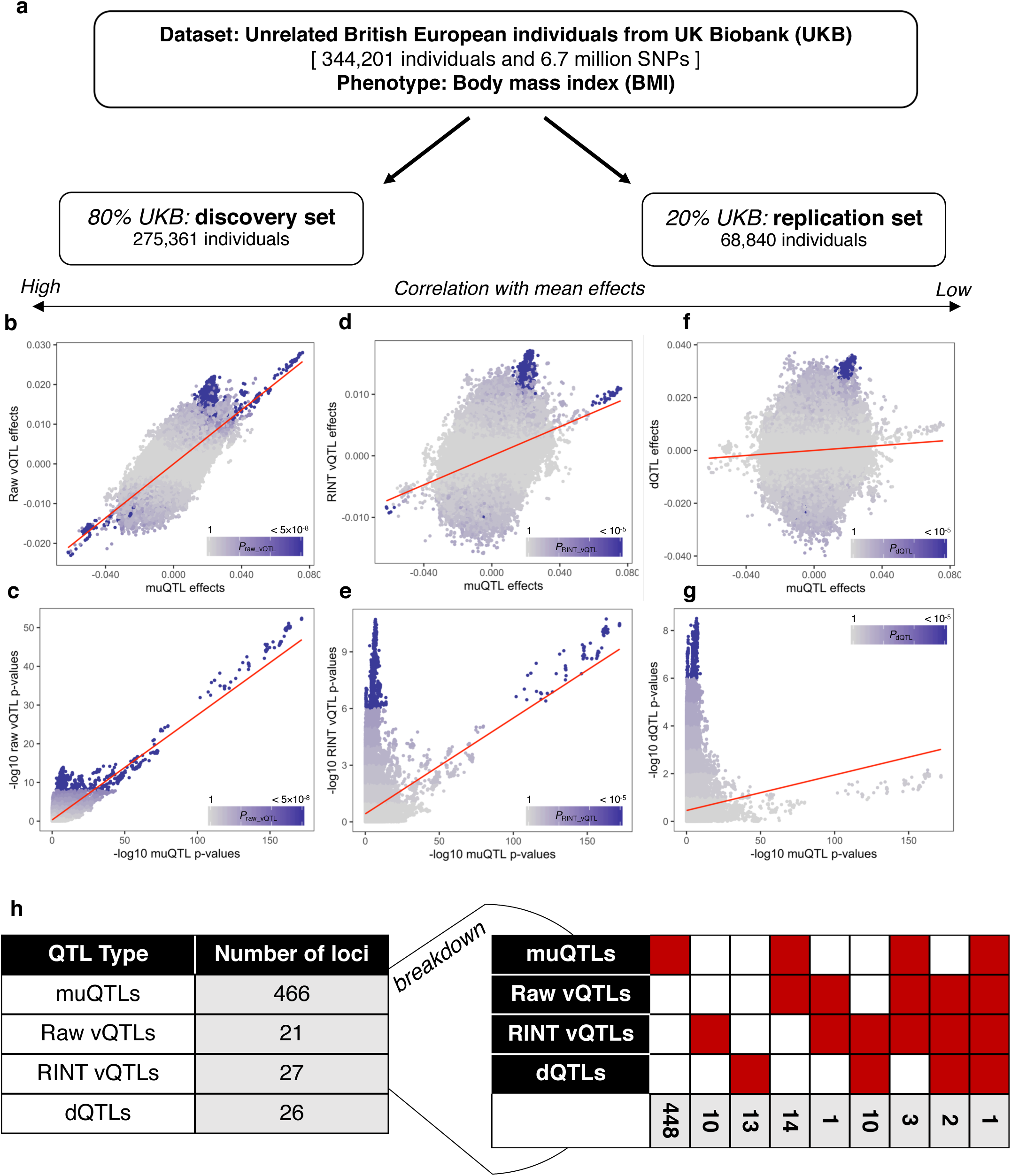
GWAS of body mass index levels in UK Biobank. (a) Data for imputed genotypes and BMI in unrelated British European individuals were split into a discovery set, representing 80% of the data, and a replication set, representing 20% of the data. Within the discovery set, a GWAS was performed on the means (muQTLs) and variances (raw vQTLs) of untransformed BMI and on the variances (RINT vQTLs) and dispersion (dQTLs) of RINT BMI. Across SNPs, the effect sizes (b) and *P*-values (c) were highly correlated between muQTLs and raw vQTLs. The RINT reduced mean-variance correlation (d) and identified a set of RINT vQTLs with smaller muQTL effects (e). Dispersion effects had the least correlation with mean effects (f), and all dQTLs were not the most significant muQTLs (g). In (b-g), the red line represents the line of best fit. Points are colored by the –log_10_ p-value of the y-axis analysis, with purple representing significant (*P* < 5 × 10^−8^ with raw BMI, *P* < 10^−5^ with RINT BMI). The GWAS results are summarized in (h), broken down into by the number of QTLs passing the different criteria (indicated by the red coloring and grey counts).

We discovered a strong correlation between mean and variance effects, which we refer to as the *mean-variance relationship* (**Fig 3b-c**). The mean-variance relationship could be explained in a number of ways. Since the sample means and sample variances are correlated in non-normal distributions and BMI is non-normally distributed, variance effects could be a consequence of a SNP’s mean effect and therefore any observed associations with phenotypic variance are not indicative of underlying interactions^25^ (Supp Note 3). Alternatively, we hypothesized that a SNP associated with the mean value of a phenotype is also more likely to be involved within interactions, thus creating the correlation we observe. Biologically, SNPs with a main effect (directly impacting the studied trait) may have a greater likelihood of having an interaction effect. Statistically, variance estimates have larger standard errors than mean estimates in a population sample; thus, interactions must have larger effect sizes to detect a change in variance and consequently mean effects would be detected too (Supp Note 4; Supp Fig 3). Therefore, a correlation between mean and variance effects might be due to both real biological and statistical causes.

To disentangle the mean-variance relationship, Young *et al*. described how analysis of a phenotype with a rank inverse normal transformation (RINT) decorrelates mean and variance effects^25^ and proposed a dispersion effect test to identify differences in variances not driven by the mean effects (known as dispersion effects). We sought to find SNPs associated with both the variance and dispersion of BMI after a rank inverse normal transformation. We used the DRM to identify variance effects and Young *et al*.’s dispersion effect test to discover dispersion effects. SNP associations were identified using a less-conservative *P* < 10^−5^ threshold to produce an expanded set of SNPs, since these analyses resulted in conservative *P*-value distribution (**Fig 3d-g**; Supp Fig 4; Supp Notes 5-6).

**Figure 4:**
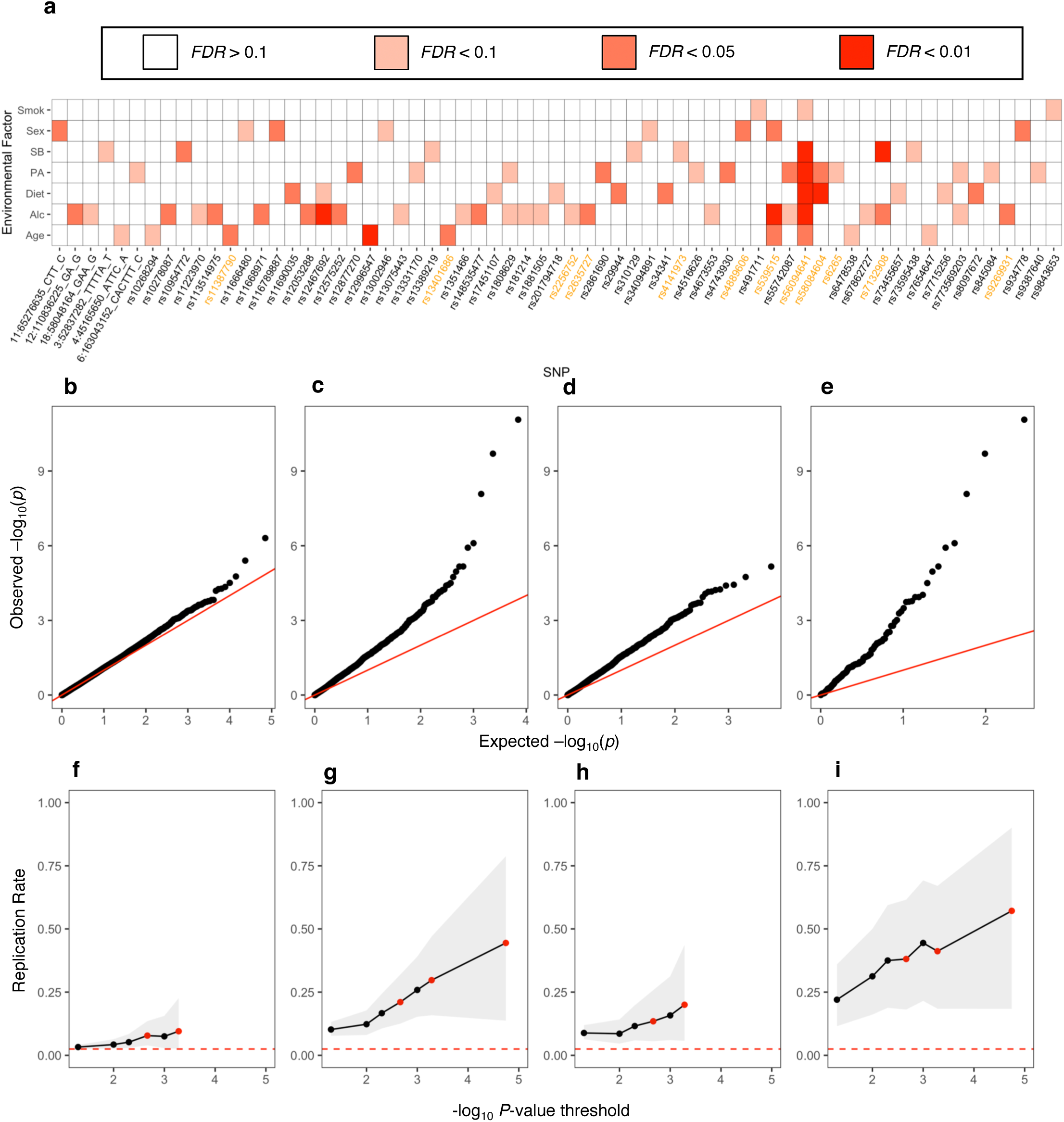
Discovery and replication of GxE interactions. (a) Heatmap of all QTLs with a *FDR* < 0.1 GxE interaction in the discovery set. Each box colored by significance level in the discovery set. Raw vQTL SNPs are highlighted in orange. Smok = smoking status; SB = sedentary behavior level; PA = physical activity level; Alc = alcohol intake frequency. (b-c) Quantile-quantile plots for all GxE interactions across environmental factors and (b) 5016 matched genome-wide SNPs, (c) 502 QTLs, (d) 448 muQTLs that are not raw vQTLs, or (e) 21 raw vQTLs. The *x*-axis shows the –log_10_ *p*-values under the null distribution and the *y*-axis shows the observed –log_10_ *p*-values, where each point represents a different GxE interaction. The red line represents the expectation under the null, with intercept = 0 and slope = 1. (f-i) Replication rates of GxE interactions, as quantified by those with the same direction of effect in both discovery and replication sets and *P*_R_ < 0.05. Given a threshold *x* (x-axis), the replication rate (y-axis) is calculated for all interactions with *P*_D_ < *x*. (f) GxE interactions using 5016 matched genome-wide SNPs. (g) GxE interactions using all 502 QTL-nominated SNPs. (h) GxE interactions using 448 muQTLs that are not raw vQTLs. (i) GxE interactions using 21 raw vQTLs. In (f-i), the confidence interval over replication rates is shown in grey and the expected replication rate under random observations (2.5%) is shown in red. Red points are *FDR* < 0.1, < 0.05, and < 0.01 cut-offs. In (g) and (i), there are no *FDR* < 0.1 associations.

We discovered 448 SNPs associated with the mean of untransformed BMI values (which we refer to as “muQTLs”), 21 SNPs associated with the variance of untransformed BMI values (“raw vQTLs”), 27 SNPs associated with the variance of transformed BMI values (“RINT vQTLs”) and 26 SNPs associated with the dispersion of transformed BMI values (“dQTLs”) (**Fig 3h**). As expected, the correlation with mean effects decreases from raw vQTLs to RINT vQTLs to dQTLs; for example, 18 of 21 raw vQTLs are also muQTLs, 4 of 27 RINT vQTLs are muQTLs, and 3 of 26 dQTLs are muQTLs. We combined the muQTLs, raw vQTLs, RINT vQTLs, and dQTLs into a set of 502 unique QTLs. We next proceeded to the second step of our sequential GxE discovery framework, where we searched for pairwise interactions between the 502 unique QTLs and with age, sex, and five environmental factors: smoking status, diet, physical activity, sedentary behavior, and alcohol intake frequency (the details of these factors are described within the Methods section).

### Discovery and replication of gene-environment interactions

To identify GxE interactions potentially explaining BMI, we used the same set of 275,361 unrelated European individuals. We tested for 3,514 GxE interactions, using the 502 unique QTLs and seven factors, by applying 3,514 distinct linear models containing a single interaction term. Overall, we identified 78 significant gene-environment interactions associated with BMI in the discovery set (*FDR* < 0.1) (**Fig 4a**), a 155-times greater discovery rate over interaction testing based on a genome-wide sampled set of SNPs (2.2% versus 0.014%; *P* = 9.8 × 10^−140^) with no expected FPR increase (**Figure 4b-c**; Methods; Supp Note 7-8; Supp Fig 5a-d).

**Figure 5:**
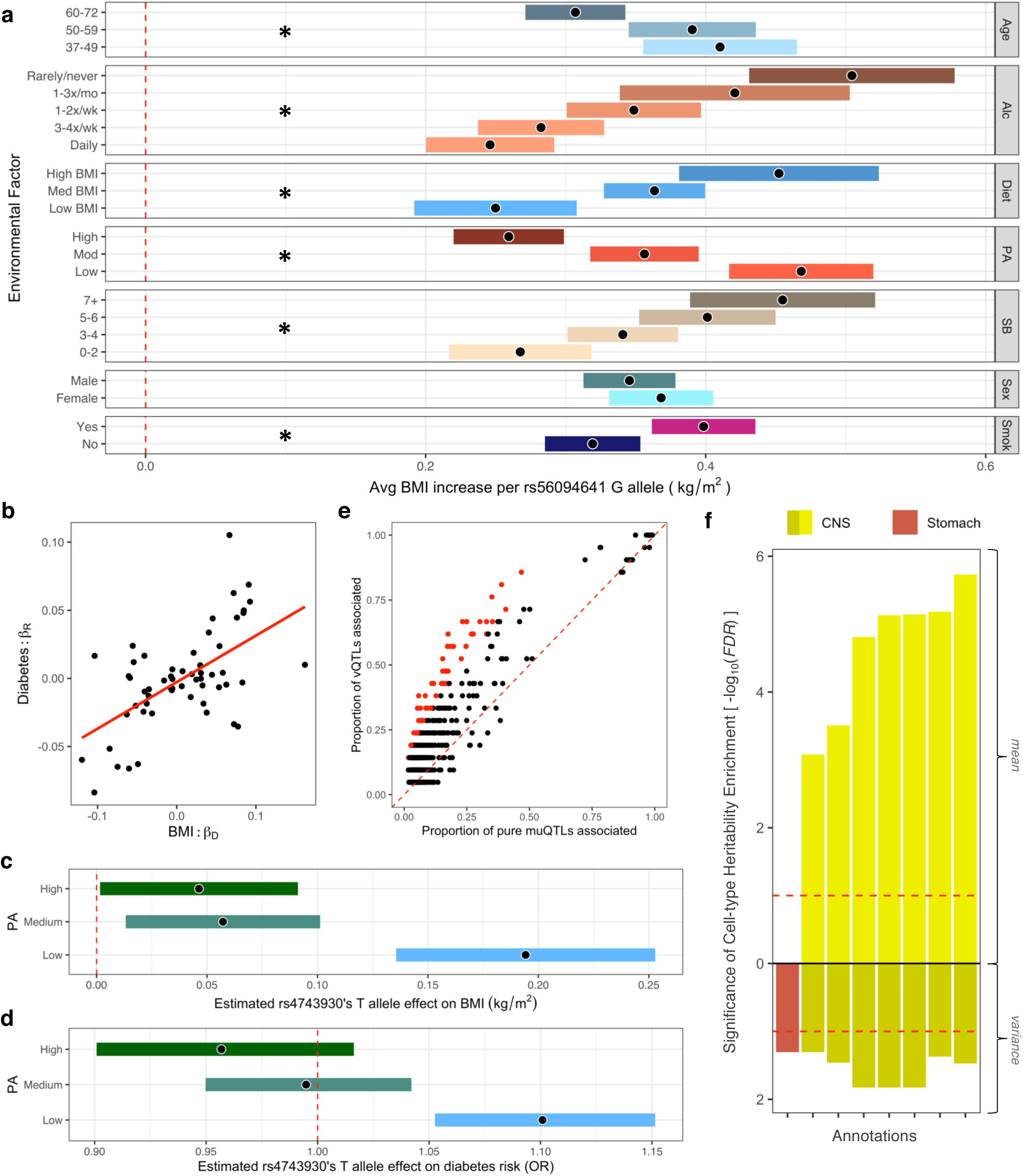
GxE interactions across environmental factors, human phenotypes, and cell types. (a) The estimated marginal BMI effect of the rs56094641 G allele conditioned on the different environmental co-variates. For visualization, age, sedentary behavior values, and diet (bottom 20%, middle 60%, upper 20%) were grouped and “rarely” or “never” answers for alcohol intake frequency were combined. Significant GxE interactions highlighted with an asterisk. (b) Estimated GxE effects in BMI within the 80% discovery set (x-axis) from linear regression were correlated with estimated GxE effects on diabetes risk within the 20% replication set (y-axis) from logistic regression. Each data point represents a different SNP x co-factor interaction. BMI GxE interactions appear predictive of diabetes GxE interactions. (c-d) The estimated marginal effect of the rs4743930 T allele on (c) BMI and (d) diabetes risk, conditioned on physical activity levels. Estimated diabetes risk effect is in terms of the relative odds ratio (OR). In (a), (c-d), the estimate is shown by the black dot, and the bars indicate the 95% confidence intervals. Smok = smoking status; SB = sedentary behavior level; PA = physical activity level; Alc = alcohol intake frequency. (e) The proportion of pure muQTLs (those with no significant raw vQTL association) associated with a phenotype were compared to the proportion of raw vQTLs that are associated. Each point is a different phenotype that is included in the Open Targets database. Phenotype associations significantly enriched in the raw vQTL set (*FDR* < 0.1) are highlighted in red. (f) The -log_10_(*FDR*) describe the partitioned enrichment of BMI mean and BMI variance heritability in specifically expressed genes for a given cell type. Only cell-types with *FDR* < 0.1 in the BMI variance analysis are shown. Dashed red lines drawn at *FDR* < 0.1.

We used an independent and randomly selected replication set of 68,840 unrelated British European individuals from UKB to evaluate our findings in the discovery cohort (**Fig 3a**). We refer to effect size estimates and *P*-values from the discovery set as β_D_ and *P*_D_, and those from the replication set as β_R_ and *P*_R_. We considered an interaction to be replicated if the direction of effect was the same in both discovery and replication sets [sign(β_D_) = sign(β_R_)] and *P*_R_ < 0.05. Overall, 21.1% of significant GxE interactions (*FDR* < 0.1) replicated, compared to 7.8% of GxE at similar nominal *P*_D_-values based on genome-wide SNPs (2.7-fold enrichment, *P* = 2.2 × 10^−4^) (**Fig 4f-g**). The estimated replication rate increased as the significance threshold became stricter, with all 9 *FDR* < 0.01 interactions replicating effect direction and 4 of the 9 passing *P*_R_ < 0.05 significance. Our data suggests that interactions using the 502 unique QTLs had significantly greater replication rates compared to interactions from genome-wide SNPs, despite similar nominal *P*_D_-values (Supp Note 9; Supp Fig 5e).

We found that the increased discovery and replication rates are driven by raw variance effects. 14.2% of tested GxE interactions using raw vQTLs were significant (*FDR* < 0.1), a 10.0-, 8.6-, and 7.7-fold higher GxE discovery rate than muQTLs, RINT vQTLs, and dQTLs in the absence of a significant raw vQTL association (1.4%, 1.7%, and 1.9% respective discovery rates, *P* < 10^−12^ for each; **Fig 4d-e**; Supp Fig 5f-i). Similarly, the interactions from vQTLs (*FDR* < 0.1) had 2.8-fold higher replication rate compared to muQTLs that were not raw vQTLs (38.1% versus 13.5%; *P* = 4.3 × 10^−3^) (**Fig 4h-i**; Supp Note 10). Lastly, we found that the GxE effects correlated best with the raw vQTL effects compared to the muQTL, RINT vQTL, or dQTL effects (Supp Fig 6). Hence, we found that the ability to discover and replicate interactions was primarily driven by a single-SNP’s marginal association with untransformed BMI variance.

### Large gene-environment interactions influence BMI

The most significant interaction identified with respect to BMI was between the *FTO* intronic region and alcohol intake frequency (*P* = 8.6 × 10^−12^). The *FTO* intron region harbors the strongest muQTL (*P* = 1.3 × 10^−172^), raw vQTL (*P* = 5.4 × 10^−53^), and RINT vQTL (*P* = 3.7 × 10^−11^) association with body mass index (rs56094641), has been functionally implicated as a key obesity regulator in mouse experiments and CRISPR-Cas9 editing of human patient samples^35,36^, and was recognized as a gene-environment interaction hotspot in previous studies^37^. At this locus, we identified additional interactions (*FDR* < 0.1) with sedentary behavior (*P*_D_ = 1.2 × 10^−6^), physical activity (*P*_D_ = 2.0 × 10^−10^), diet (*P*_D_ = 7.9 × 10^−7^), age (*P*_D_ = 1.1 × 10^−4^), and smoking behavior (*P*_D_ = 9.6 × 10^−4^), but not with sex (*P*_D_ = 0.39) (**Table 1** and **Fig 5a**). We fit a model containing each significant interaction with rs56094641 and found a significant effect for each GxE term (*P* < 0.05), suggesting that each interaction is independent (Supp Table 1).

**Table 1:** GxE interactions with *FDR* < 0.01. From left to right: The SNP name, annotated gene (based on evidence in the Open Targets database, see Methods), environmental factor (Smok = smoking status; SB = sedentary behavior level; PA = physical activity level; Alc = alcohol intake frequency), estimated effect size and *P*-values in the discovery cohort, estimated effect size and *P*-values in the replication cohort, and *P*-values from the four QTL studies: muQTLs, raw vQTLs, RINT vQTLs, and dQTLs (colored in red if significant).

| SNP | Gene | E | $\beta_D$ | $P_D$ | $\beta_R$ | $P_R$ | $P_{\text{mean}}$ | $P_{\text{raw}}$ | $P_{\text{RINT}}$ | $P_{\text{disp}}$ |
| --- | --- | --- | --- | --- | --- | --- | --- | --- | --- | --- |
| rs539515 | SEC16B | Alc | -0.045 | $1.1 \times 10^{-5}$ | -0.065 | $1.6 \times 10^{-3}$ | $10^{-44}$ | $10^{-8}$ | 0.26 | 0.42 |
| rs56094641 | FTO | Alc | -0.058 | $8.6 \times 10^{-12}$ | -0.072 | $2.1 \times 10^{-5}$ | $10^{-172}$ | $10^{-53}$ | $10^{-11}$ | 0.01 |
| rs56094641 | FTO | SB | 0.025 | $2.5 \times 10^{-6}$ | 0.011 | 0.26 | $10^{-172}$ | $10^{-53}$ | $10^{-11}$ | 0.01 |
| rs56094641 | FTO | PA | -0.103 | $2.0 \times 10^{-10}$ | -0.077 | 0.02 | $10^{-172}$ | $10^{-53}$ | $10^{-11}$ | 0.01 |
| rs56094641 | FTO | Diet | 0.078 | $9.6 \times 10^{-8}$ | 0.091 | $3.5 \times 10^{-4}$ | $10^{-172}$ | $10^{-53}$ | $10^{-11}$ | 0.01 |
| rs58084604 | MC4R | Diet | 0.084 | $7.2 \times 10^{-7}$ | 0.049 | 0.10 | $10^{-73}$ | $10^{-19}$ | $10^{-4}$ | 0.21 |
| rs7132908 | FAIM2 | SB | 0.030 | $8.4 \times 10^{-9}$ | 0.008 | 0.44 | $10^{-30}$ | $10^{-11}$ | $10^{-3}$ | 0.14 |
| rs12467692 | UBE2E3 | Alc | 0.040 | $6.8 \times 10^{-6}$ | 0.031 | 0.08 | $10^{-10}$ | $10^{-3}$ | 0.26 | 0.96 |
| rs12996547 | TMEM18 | Age | -0.007 | $1.8 \times 10^{-5}$ | -0.006 | 0.07 | $10^{-16}$ | $10^{-7}$ | $10^{-3}$ | 0.05 |

The most significant GxE interaction identified outside the *FTO* region was between the rs7132908 variant and sedentary behavior level (*P*_D_ = 8.4 × 10^−9^). This variant lies in the 3’-UTR of the *FAIM2* gene, with selective and functional constraint (as estimated by SiPhy^38^ and GERP^39^), 30 different bound proteins in ENCODE ChIP-Seq experiments^40^, and a high prevalence of enhancer histone marks and DNAse sites across tissue types in the Roadmap Epigenomics Consortium^41,42^. *FAIM2* encodes an anti-apoptopic protein and exhibits a brain-specific gene expression pattern across human tissues^43^. Previously, it has been shown that *FAIM2* gene expression is regulated by dietary exposures^44^ and *FAIM2* promotor methylation is regulated by sedentary behavior^45^. Furthermore, the rs7132908 variant has another significant GxE interaction (*FDR* < 0.1) with alcohol intake frequency on BMI (*P*_D_ = 1.7 × 10^−4^).

The four other *FDR* < 0.01 gene-environment interactions we discovered were between rs58084604 (near *MC4R*) and diet (*P*_D_ = 7.2 × 10^−7^), rs539515 (*SEC16B*) or rs12467692 (*UBE2E3*) and alcohol intake frequency (*P*_D_ = 1.1 × 10^−5^ and *P*_D_ = 6.8 × 10^−6^ respectively), and rs12996547 (*TMEM18*) and age (*P*_D_ = 1.8 × 10^−5^). All 9 *FDR* < 0.01 interactions replicated with the same direction of effect, and 4 of the 9 had *P*_R_ < 0.05 (**Table 1**). *MC4R* gain-of-function mutations protect against obesity risk^46^ and have been functionally validated in obesity within mice^47^, while *Sec16b* knockout mice carry decreased cholesterol levels with higher body weight^48^. We found that age and the rs12996547 haplotype are associated with increased *TMEM18* gene expression in visceral adipose GTEx tissue^43^ (age: *r* = 0.22; *P* = 4.4 × 10^−5^; SNP: *β* = 0.155, *P* = 3 × 10^−4^), which may be one mechanism to jointly reduce BMI levels. Previously, *Tmem18* germline loss in mice led to increased body weight, while over-expression resulted in weight loss by regulating appetite and energy balance^49^. Our findings in UKB and GTEx lend further evidence to support the role of *TMEM18* in BMI.

### GxE interactions have pleiotropic effects over BMI and diabetes risk

We aimed to determine whether gene-environment interactions influencing body mass index levels exhibit pleiotropic effects and are shared across human diseases, possibly by jointly influencing BMI and disease risk (Supp Fig 7a). From the set of 78 significant GxE interactions above (*FDR* < 0.1), we identified a set of 58 GxE interactions associated with the same direction of effect on BMI in both discovery and replication sets. We then screened these GxE interactions against three additional medical diagnoses: coronary artery disease, diabetes, and high blood pressure diagnosis (Methods).

We found that GxE effects on BMI estimated in the discovery cohort significantly correlated with GxE effects on diabetes risk within the held-out set (*r* = 0.59, *P* = 1.3 × 10^−6^) (**Fig 5b**), even after adjusting for BMI as a confounder (*r* = 0.38, *P* = 3.3 × 10^−3^) (Supp Fig 7b), indicating that BMI GxE effects are predictive of GxE influences over diabetes risk. Furthermore, we identified one significant disease interaction (*FDR* < 0.1), where physical activity regulated the association of rs4743930 with diabetes risk (*P* = 8.7 × 10^−5^; *FDR* = 0.015). In a previous UKB analysis^50^, this variant is marginally associated with diabetes risk at *P* = 0.022, and consequently would not appear as one of the most significant findings in a hypothesis-free GWAS. Within low exercise individuals, the rs4743930 T allele was associated with increased BMI levels (β_D_ = 0.19 kg/m^2^ per T, *P*_D_ = 8.5 × 10^−11^) and increased diabetes risk (*OR*_D_ = 1.10, or a 10% risk increase per T, *P*_D_ = 2.7 × 10^−5^). Within moderate or high exercise individuals, there was a minor association with BMI levels and no significant association with diabetes risk (**Figure 5c-d**). This interaction could be linked to decreased BMI levels and protective diabetes effects in both discovery and replication sets (BMI: β_D_ = -0.075, β_R_ = -0.042; diabetes risk: β_D_ = - 0.075, β_R_ = -0.065), although *P*_R_ = 0.25 and *P*_R_ = 0.09 for BMI and diabetes risk in the replication set (possibly due to lower sample size, as Table 1 shows that more than half of the *FDR* < 0.01 interactions had *P*_R_ > 0.05 despite same effect direction). The observed associations remained present after adjusting for BMI as a confounder (β_D_ = -0.058, *P*_D_ = 2.5 × 10^−3^; β_R_ = -0.066, *P*_R_ = 0.09) (Supp Fig 7c).

Leveraging the genomic (transcription start site proximity), transcriptomic (eQTL studies), and epigenomic information (Promotor Capture Hi-C data) found in the Open Targets database^51^, we inferred that rs4743930 likely regulates the *BARX1* gene, which is part of the homeobox transcription factor family integral to anatomical development. *BARX1* exhibits a noteworthy tissue-specific gene expression pattern across human tissues, with high expression in visceral adipose, esophagus, and stomach tissue and very low expression in other GTEx tissues^43^ (Supp Fig 7d). Previous research has shown that the Barx1 transcription factor protein is a key regulator of stomach cell fate and organogenesis and *Barx1*^*-/-*^ knockout mice have significantly altered stomach morphology due to inhibition of the Wnt signaling pathway^52,53^. As the Wnt signaling pathway modulates the formation of adipose tissue and regulates the sensitivity to insulin, it has been proposed that pathway malfunctioning could lead to high co-morbidities of obesity and diabetes^54^. Here, we provide novel human genetic evidence of a pathway regulator, *BARX1*, to support Wnt signaling’s proposed pleiotropy over body weight and diabetes risk.

### Evidence for weak epistatic interactions associated with BMI

While the primary goal of this study was the discovery of GxE interactions, we hypothesized that a similar approach could be used to discover gene-gene (GxG) interactions in relation to BMI. We first tested for GxG interactions associated with BMI levels by performing all-pairwise interaction testing between 502 QTLs (125,751 tests). We found no departure from the -log_10_(*p*-values) expected under the null distribution and there was no correlation between interaction effects estimated in the 80% discovery cohort versus 20% replication cohort (r = - 0.003, *P* = 0.30) (Supp Fig 8a-b). Most importantly, unlike GxE interactions, leveraging mean, variance, or dispersion effects did not provide a reliable inroad to discovering GxG interactions (Supp Note 11; Supp Fig 8e). However, when considering the more significant interactions (*P*_D_ < 0.001), we observed a weak correlation between effects estimated in each cohort (*r* = 0.17, *P* = 0.04) and found that statistical replication rates increased slightly above the theoretical null, 2.5- 3% (Supp Fig 8c-d). Our results in BMI suggest that any potential underlying epistatic effects are small and would be difficult to detect, concordant with a recent search for epistasis in three biologically simpler molecular traits^55^.

### vQTLs are linked to environmentally-influenced pathways and phenotypes

Since SNPs associated with the variance of untransformed BMI acted as hotspots of GxE interactions, we explored whether certain phenotypes and pathways were more likely to be linked to raw vQTLs compared to SNPs only associated with the mean of BMI. These muQTLs which are not significant vQTLs are referred to as “pure muQTLs”.

To evaluate this, we used the Open Targets database^51^ which contains a large catalog of genotype-phenotype associations. We performed a (mean-based) phenotype-wide association study (PheWAS) of 21 raw vQTLs and 448 pure muQTLs by querying all phenotypes available in Open Targets. Using a binomial test, we assessed whether the group of raw vQTLs were enriched for an association with a phenotype (nominal *P* < 0.05) compared to the group of pure muQTLs (nominal *P* < 0.05) (Methods).

Overall, we found vQTLs were enriched for an association with many phenotypes that have a strong environmental influence (whether from diet, exercise, infection, or microbiome). These included several diabetes-, immune-, and hematological-related phenotypes (**Table 2, Fig 5e**). Permutation analyses of the pure muQTLs showed that the PheWAS-based enrichment test did not have inflation of false positives (Supp Fig 9a).

**Table 2:** PheWAS enrichment of raw vQTLs versus pure muQTLs. From left to right: the phenotype, the proportion of pure muQTLs and raw vQTLs that are associated with the phenotype, the ratio between the two proportions, the binomial test *P*-value to assess vQTL set enrichment, and the *FDR* corrected significance. These phenotypes represent a manually-curated and incomplete list of all significant findings presented in Supp Data 9.

| Phenotypes | muQTL | vQTL | Ratio | *P* | *FDR* |
| --- | --- | --- | --- | --- | --- |
| Diabetes diagnosed by doctor | 0.47 | 0.86 | 1.83 | 2.64E-04 | 0.017 |
| Diabetes mellitus | 0.33 | 0.67 | 2.029 | 1.56E-03 | 0.047 |
| Diabetic retinopathy | 0.05 | 0.33 | 6.318 | 6.85E-05 | 0.010 |
| Eosinophil counts | 0.15 | 0.43 | 2.882 | 1.89E-03 | 0.055 |
| Hypothyroidism | 0.12 | 0.43 | 3.647 | 3.32E-04 | 0.020 |
| Mean corpuscular haemoglobin | 0.23 | 0.67 | 2.866 | 2.87E-05 | 0.009 |
| Neutrophil percentage | 0.23 | 0.52 | 2.299 | 2.98E-03 | 0.065 |
| Osteoarthritis I non-cancer illness code, self-reported | 0.17 | 0.57 | 3.310 | 4.36E-05 | 0.010 |
| Red blood cell (erythrocyte) distribution width | 0.27 | 0.62 | 2.305 | 7.89E-04 | 0.038 |
| Red blood cell count | 0.20 | 0.57 | 2.906 | 1.64E-04 | 0.014 |
| Reticulocyte fraction of red cells | 0.18 | 0.62 | 3.536 | 7.18E-06 | 0.006 |
| Type 1 diabetes | 0.09 | 0.33 | 3.861 | 1.40E-03 | 0.047 |
| Type 2 diabetes | 0.41 | 0.71 | 1.762 | 4.13E-03 | 0.078 |
| Type 2 diabetes with neurological manifestations | 0.06 | 0.38 | 6.619 | 1.24E-05 | 0.006 |
| Type 2 diabetes with ophthalmic manifestations | 0.06 | 0.33 | 5.148 | 2.47E-04 | 0.017 |
| Ulcerative colitis I non-cancer illness code, self-reported | 0.05 | 0.24 | 4.513 | 4.08E-03 | 0.078 |

Next, we mapped non-MHC SNPs to single genes using genomic proximity and Open Targets’ variant-to-gene pipeline, queried raw vQTL gene sets or pure muQTL gene sets in GeneMania, and performed gene ontology (GO) enrichment analysis of the resulting gene network (Methods; Supp Note 12; Supp Data 1-2). We found that the network of raw vQTL genes was enriched for G protein coupled receptor-related (GPCR) signaling pathways and cell growth processes (Supp Data 3-4). In contrast, the pure muQTL network was enriched for developmental processes, particularly in the central nervous system (CNS) with no enrichment in the GPCR-related GOs (Supp Data 5-6). GPCRs transduce extracellular signals and activate downstream a cascade of intracellular proteins and pathways, which is essential for how cells interact with the environment.

### Polygenic heritability analysis implicates stomach cell types in regulating BMI variance

We evaluated whether the genetic contribution to the variance of BMI, a potential proxy for GxE interactions, might implicate different cell types in regulating BMI levels compared to studies on the mean of BMI. In Drosophila, the variance of a phenotype can be heritable^56^. We performed partitioned linkage disequilibrium score regression on mean and variance GWAS summary statistics to find cell types enriched for mean or variance heritability. We used 205 functional annotations from GTEx^43^ and the Franke lab^57^ that describe tissue-specific genes in each cell type.

Overall, we found that estimated cell type enrichment values were similar for BMI means and variances (*r* = 0.81; *P* = 1.4 × 10^−48^) (Supp Fig 9b-c). For example, the genetic signal for both means and variances were clustered in genes uniquely expressed in the CNS, as described previously^58^. Notably, we discovered that the heritability of BMI variance was significantly enriched (*FDR* < 0.1) at genes with the highest expression in stomach cell types (*P* = 1.2 × 10^−3^; *FDR* = 0.049), with no significant association in these regions for mean heritability (*P* = 0.40) (**Fig 5f**; Supp Note 13). This preliminarily suggests that stomach cell types, in addition to CNS cell types, have a critical role over BMI variance and regulating potential GxE interactions, and that this would not be discovered in a mean-based analysis.

## Discussion

We have identified SNPs associated with the variance of BMI (vQTLs), which are enriched for gene-environment (GxE) interactions and for associations with phenotypes under strong environmental influences. When functionally profiling the annotated genes of vQTLs, we found enrichment for G protein coupled receptor-related signaling pathways, which are key to cells’ responses to the external environment. We also discovered through a polygenic analysis that a significant proportion of the heritability in BMI variance is clustered near genes highly expressed in stomach cells, which were not revealed in an analysis of BMI means^58^. Future application of our methods across phenotypes has the potential to identify genes, pathways, or cell types that serve as key regulators of the interplay between genetics and environment.

Additionally, we showed how GxE interactions first identified in BMI were predictive of the GxE effects on diabetes risk within a distinct set of individuals. We further discovered a *BARX1* regulatory locus that significantly increases BMI and diabetes risk in low exercise individuals but does not have pleiotropic population effects in moderate to high exercise individuals. This framework of screening for SNPs as interaction candidates within quantitative phenotypes to subsequently discover interactions influencing complex disease can be broadly applicable across the range of human phenotypes. Methods to deconvolute case-control disease phenotypes into a quantitative scale that re-captures disease granularity and severity will enable the application of vQTL testing directly to the disease phenotype of interest.

We explored multiple approaches to decouple mean and variance effects, evaluate the relationship between the two, and find GxE interactions. While Young *et al*.^25^ introduced a test for identifying SNPs associated with the variance of a phenotype independent of a mean effect (which we referred to as dQTLs), we found that the strongest GxE signal came from the SNPs associated with the variance of BMI prior to statistical transformations (raw vQTLs). If raw vQTLs are a robust footprint of interactions and estimated raw vQTL effects correlated strongly with mean-based effects, then this suggests that any SNP directly impacting BMI may be more likely to have its BMI effect modified by another factor.

GWAS-type testing is not the only approach to limiting the number of potential interactions to explore. Other previously used approaches for reducing vast genomic data are to filter SNPs based on prior biological information^59^ or to combine SNPs into higher-order gene-level data^60^. Alternatively, a large number of environmental variables can be combined into a single environmental score^61^. A significant drawback of these methods is that they will not be a hypothesis-free genome-wide approach to discover the epidemiological interaction between SNPs and other factors. Prior information is biased to prior knowledge, and gene-level data or an environmental score limits the search space for potential interactions.

One future research area is the evaluation of polygenic scores that consider interaction effects. Polygenic scores are currently based on only marginal additive effects, and our research identified strong GxE interactions influencing BMI variability. For example, variants in the *FTO* intron region (the strongest genetic regulators of obesity) are associated with a nearly double BMI increase in low exercise individuals compared to high exercise individuals (**Fig 5a**). Interactions can perturb each individual from the expectation given a single genotype, and the ideal individual prediction would accommodate these interaction effects.

While the DRM vQTL approach that was applied in these analyses has advantages in power, any detection of interaction effects will have lower power than tests of main effects. It is anticipated that the increasing sample sizes of GWAS will enable more sensitive detection of significant loci and more precise estimation of their variance effects. This will in turn improve the sensitivity of a variance test in detecting underlying GxE interactions. Furthermore, replication of GxE interactions require special attention. Here we split UK Biobank individuals into two mutually exclusive groups, but this approach is not the same as performing tests on two completely independent population samples. There may be unmeasured confounding factors in the UK Biobank samples that drive spurious associations. Interactions will need to be independently replicated in other cohorts to weed out spurious signals, although any lack of replication could be due to differences in allele frequencies, cultural behavior, and other environmental variables. For our purposes, we used the study design to allow a comparison of replication rates between two sets of GxE interactions.

Perhaps the most important requisite to improve our understanding of GxE interaction in humans is the collection of accurate, high-quality measurements of relevant environmental variables. Specialized wearable tracking devices and improvements in biomarker data are being explored, and the hope is that these will deliver a quantum improvement in the availability and accuracy of environmental data. In these settings, vQTLs can provide a promising approach to reduce dimensionality of genetic data and increase statistical power to detect GxE interactions. Overall, our work highlights the ability to discover significant environmental influences that modulate the genetic contribution to human phenotypes.

## Methods

### Description and implementation of the variance tests

In the DRM, each individual *j* with genotype *i* has phenotypic value *Y*_*ij*_. The genotype is coded as *i* = 0, 1, or 2, determined by the minor allele count. The median phenotype value is calculated for all individuals with categorical genotype *i*, 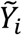. The absolute value of the difference between *Y*_*ij*_ and 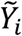 is calculated:

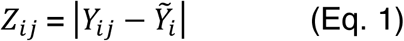

The *Z*_*ij*_ values for each individual *j* represent the deviation from the within-genotype phenotype medians. Next, SNPs are tested for association with *Z*_*ij*_ values using linear regression and the genotype indices as a numeric covariate. The effect size and *P-*values for the SNP covariate in the regression are used as proxies for the variance effect size and significance of association with phenotypic variance. The DRM is a similar approach to the Levene’s test, which allows for non-linear associations through an ANOVA model instead of a linear model. In practice, covariates are regressed out from *Y*_*ij*_ prior to calculating *Z*_*ij*_.

In our study, we used a number of other variance tests. The two-step squared residual approach was implemented as linear regression on the squared mean-centered phenotype. The other variance tests were implemented in *R* using the *dglm()* function from the *dglm* package, the *bartlett*.*test()* and *fligner*.*test()* function from the *stats* package, and the *leveneTest()* function with default arguments (median-centered) from the *car*^*62*^ package.

### Simulations of genotypes and phenotypes for method comparison

The methods for identifying variance differences were compared using statistical power and false positive rates (FPR) as the performance benchmarks. In the FPR scenario, a single SNP was simulated with MAF = 0.4 using a *W* ∼ *Binom*(2, 0.4) independent random variable. The SNP value *W* is set to be equal to the genetic component, *Y*_G_, of a phenotype. In the power testing scenario, a SNP *X*_1_ and an environmental factor *X*_2_ were simulated using a binomial distribution with probability of success = 0.4: *X*_*1*_ ∼ *Binom*(2, 0.4) and *X*_*2*_ ∼ *Binom*(2, 0.4). This other factor can also be thought of as an environmental exposure with three levels (e.g., for physical activity: never exercise, rarely exercise, often exercise). The product *X*_*1*_ × *X*_*2*_ is the genetic component, *Y*_G_, of a phenotype in the power scenario. When contrasting muQTL approaches to vQTL approaches, the genetic component is equal to the product *X*_*1*_ × (*X*_*2*_ – 1). This is so that *X*_*1*_’s marginal association with the phenotype is either positive or negative, depending on the value of *X*_2_.

Phenotypes were generated by summing a point genetic effect, *Y*_G_, and random environmental noise, *Y*_E_. *Y*_E_ was simulated from a normal distribution or a chi-squared distribution with 4 degrees of freedom, and scaled appropriately such that *Y*_G_ explains *Ψ* proportion of the variance in the phenotype and *Y*_E_ explains 1 – *Ψ* percent, as described below.

Given *Ψ*, the proportion of the variation explained by the environmental component is larger than *Ψ* by a factor 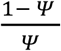. After calculating the variance of the genetic component, *V*_G_, the variance of the environmental noise, *V*_E_, can be calculated as:

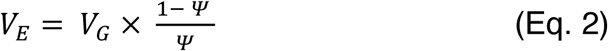

In practice, the normally distributed environmental noise can be simulated as *Y*_E_ ∼ *N*(0, *V*_E_) for normally distributed phenotypes. Chi-square distributed noise can be simulated as the following for chi-square distributed phenotypes:

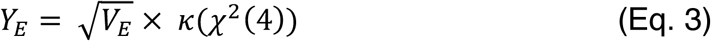

κ is the function that centers and scales the chi-square input to have mean equal to 0 and variance equal to 1. The final phenotypic values were created by calculating the sum *Y*_G_ + *Y*_E_. In all, genotypes and phenotypes were generated for 10,000 individuals.

The association between SNP and the variance in phenotype was tested using the different variance methods. The null hypothesis (no association) was rejected when the nominal *P* < 0.05. This was repeated across 1,000 simulations with distinct genotypes and phenotypes. Power and FPR refer to the proportion of simulations where the null hypothesis was rejected. LT and DRM effects were compared by analyzing the simulations where the interaction explained less than or equal to 5% variance in the phenotype.

Linear regression was used to compare a muQTL approach to a vQTL approach (using the DRM). A contingency table was calculated from count data across simulations that describes whether the muQTL method rejected the null hypothesis of no association or the vQTL method rejected the null hypothesis of no association. A two-sided Fisher exact test was performed for each variance explained value, *V*_G_, separately to assess the relationship of vQTL power with muQTL power.

### UK Biobank data

UKB data was processed previously by the UK Biobank team^63^ and accessed under Application ID 47137. The individuals and SNPs used in analysis were limited to those in the Neale lab’s analysis^50^ as the same quality control criteria were adopted for sample and genotypes in this analysis. By doing so, individuals were removed based on whether they were not used in the UKB team’s principal component analysis (removing related samples), not of European British ancestry, or had sex chromosome aneuploidy, excess heterozygosity, or outlier genotype missing rates. Genotypes were removed if INFO score < 0.8, MAF < 0.05, or HWE *P* < 10^−10^. The full processed and quality-controlled data contained 344,201 individuals and 6,701,215 SNPs.

Analysis was randomly split into two parts. A discovery set contained 80% of the data, selected randomly from the full dataset, which was used for discovering associations between SNPs and phenotypes. A replication set contained the remaining 20% of the data, which was used for the replication of associations identified in the discovery set.

### Genome-wide association study in UK Biobank

A genome-wide association study (GWAS) was performed within the discovery set containing 80% of the data. Individuals with body mass index levels greater than 5 standard deviations from the mean removed from analysis to prevent a large influence from outliers which could be driven by non-modeled factors. Body mass index levels were adjusted for the following covariates: sex, age, age × sex, age^2^, age^2^ × sex, genotyping array, and principal components 1 – 20. This was performed by fitting a linear model and calculating the residuals. Using the residuals, we performed a GWAS by using linear regression (mean effects) and the DRM (variance effects) between a single SNP and adjusted, untransformed BMI. The findings from these analyses were referred to, respectively, as muQTLs and raw vQTLs. We also applied a rank inverse normal transformation (RINT) to the residuals to decorrelate mean and variance effects, and proceeded with a GWAS using the DRM and a dispersion effect test^25^ (DET). The rank inverse normal transformation uses the ranks of phenotype values and inverse transforms the ranks into a normal distribution. We refer to the DRM and DET outcome as RINT vQTLs and dQTLs.

All genome-wide association analyses were implemented on sets of 5000 SNPs and performed in parallel. Genome-wide linear regression was performed using *plink*^64^ and the DRM was performed by employing the *BEDMatrix* R package (https://github.com/QuantGen/BEDMatrix). The DET was implemented by first using a Python-implemented heteroskedastic linear model^25^ (https://github.com/AlexTISYoung/hlmm). The dispersion effects were then estimated by using the additive and log-linear variance effects as described previously^25^; this method is implemented in the *estimate_dispersion_effects*.*R* file in the linked *hlmm* repository.

Results from these analyses were compared using correlations. Significance was determined with the criterion *P* < 5 × 10^−8^ for untransformed analyses and *P* < 1.0 × 10^−5^ for RINT results. Significant QTLs were used as the nominated loci for identifying GxG and GxE interactions. Previous GWAS results were downloaded from the Neale Lab webpage^50^.

### Construction of a diet score

We computed a diet score to be used as an interaction factor in GxE analysis by adapting a protocol described previously^25,37^. First, we extracted 18 diet-related variables: “Cooked vegetable intake”, “Salad / raw vegetable intake”, “Fresh fruit intake”, “Dried fruit intake”, “Bread intake”, “Cereal intake”, “Tea intake”, “Coffee intake”, “Water intake”, “Oily fish intake”, “Non-oily fish intake”, “Processed meat intake”, “Poultry intake”, “Beef intake”, “Lamb/mutton intake”, “Pork intake”, “Cheese intake”, and “Salt added to food”. We next fit a linear model using baseline model covariates plus the 18 diet variables. These baseline model covariates included age, sex, age^2^, age x sex, age^2^ x sex, genotyping array, and principal components 1-20^25^. We fit a model to 25% of the UK Biobank discovery set (thus, 20% of the full data set used in the study) (*N* = 68,840), and estimate β coefficients for each diet variable. In the remaining 275,361 individuals (which include those from both the discovery and replication sets), we used the estimated β coefficients for each diet variable to calculate a diet score:

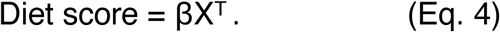

Above, β is the 1 × 18 vector of coefficients for diet variables and X is a 275,361 × 18 matrix of diet variable values for the 275,361 individuals.

Low diet score values describe a diet predicted to be associated with low BMI individuals, while high diet score values describe a diet predicted to be associated with high BMI individuals. Potential interactions with genetic polymorphisms may describe a change in the average relationship between diet score and BMI within the general population. This would suggest that the effects of the different diet variables on BMI is synergistically higher or lower than expected.

### Making non-diet environmental variables

We used UKB fields 21022-0.0, 22001-0.0, and 1558-0.0 for age, sex, and alcohol intake frequency. The alcohol intake frequency field was re-coded in the opposite direction, such that a higher value indicates a higher alcohol intake frequency. Individuals with missing data or preferred not to answer were removed. For smoking status, physical activity level (PA), and sedentary behavior level (SB), we generated new variables using the methods described in Wang *et al*.^26^.

For smoking status, we used fields 1239-0.0 (“Current tobacco smoking”) and 1249-0.0 (“Past tobacco smoking”) to create a binary variable. Individuals were only coded as 0 if they do not currently smoke, and they answer regarding their past history, “I have never smoked” or “Just tried once or twice”. Individuals were classified as 1 if they currently or previously smoke most days or occasionally. Individuals with missing data and who could not fill the criteria were removed.

For PA, we used fields 864-0.0 (“Number of days/week walked 10+ minutes”, 874-0.0 (“Duration of walks”), 884-0.0 (“Number of days/week of moderate physical activity 10+ minutes”), 894-0.0 (“Duration of moderate activity”), 904-0.0 (“Number of days/week of vigorous physical activity 10+ minutes”), 914-0.0 (“Duration of vigorous activity”), which we labeled DayW, DurW, DayM, DurM, DayV, and DurV. According to the International Physical Activity Questionnaire analysis guideline^65^, the total metabolic equivalent minutes (METT) can be approximated as:

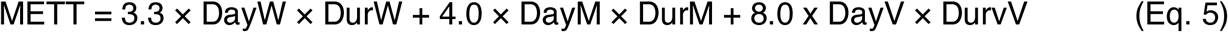

Next, PA for each individual was assigned 1, 2, or 3 for low, medium, or high activity as such:

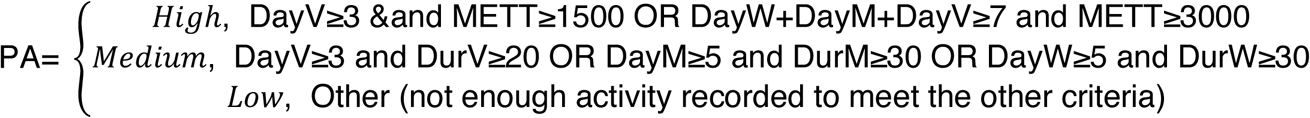

For SB, we used fields 1090-0.0 (“Time spent driving”), 1080-0.0 (“Time spent using computer”, and 1070-0.0 (“Time spent watching television (TV)”). For each variable, “Less than an hour a day” (−10) was set equal to 0 and “Do not know” or “Prefer not to answer” (−1 or -3) answers were imputed with the median of the remaining values. SB was set equal to the sum of the three columns. Outlier individuals were removed, as defined by those with greater than 5 standard deviations from the mean.

### Sampling random SNPs matched to QTLs

We calculated the frequency of homozygous minor genotypes (*f*_minor_), the minor allele frequency (MAF), and the count of individuals with a non-missing genotype at each SNP (*N*_miss_). To identify the underlying null distribution of various statistics in our study, we sample 10 matched SNPs for each QTL. Each matched SNP must have a MAF and *f*_minor_ that is below a 1% margin from the QTL, an *N*_miss_ within 1% of the QTL’s *N*_miss_ count, and be on a different chromosome than the QTL.

### Identification of genetic interactions

Pairwise interaction testing was performed between all SNP candidates and with each of the seven environmental factors in the discovery and replication sets separately. For GxE interactions with diet, only 75% of the discovery set (60% of full UKB set) was used for association tests since 25% was used to fit the model for calculating the diet score variable. GxE *P*-values were adjusted using false discovery rate and significance determined by *FDR* < 0.1. GxG *P*-values were separately adjusted using false discovery rate and significance determined by *FDR* < 0.1.

GxE discovery rate was compared between the QTL set and the genome-wide matched SNP set. First, the same number of SNPs as the QTL set were sampled from the matched SNP set. Second, the GxE *P*-values from the sampled genome-wide SNPs were adjusted using false discovery rate. Third, the discovery rate within the set was calculated as the proportion of GxE interactions with *FDR* < 0.1. This was repeated across 10,000 iterations and the mean discovery rate across the iterations was used as the expected probability in a one-sided binomial test.

### Statistical replication of genetic interactions

Genetic interactions discovered in the discovery cohort were tested for in the replication set. Given a -log_10_(*P*) threshold equal to *x*, all more significant interactions (those with -log_10_(*P*) > *x*) were identified. Within the replication cohort, an interaction is considered to have been replicated if the direction of effect was the same as in the discovery set and if the P-value in the replication set is *P* < 0.05. The replication rate is the proportion of interactions to have replicated according to these two criteria. The genome-wide replication rate was computed by using the matched and randomly sampled SNPs and testing for GxE interactions within both the 80% and 20% cohorts. The replication rate was calculated at *x* = [0.05, 0.01, 2.16 × 10^−3^, 0.005, 5.22 × 10^−4^, 1.80 × 10^−5^], where *x* = [2.16 × 10^−3^, 5.22 × 10^−4^, 1.80 × 10^−5^] are *FDR* < [0.1, 0.05, 0.01] thresholds; these were calculated by identifying the maximum GxE *P*_D_ within the QTL set that pass the respective *FDR* thresholds. In displays, the grey confidence intervals are derived from a binomial test with rate equal to 0.025, which is the theoretical replication rate under no true association.

To test for differences in replication rate, a background set is specified and the replication rate within this background set is used as the theoretical success rate in a one-sided exact binomial test. The number of successes and number of trials is from the number of replicated GxE interactions and total number of GxE interactions in the other GxE set.

### Analyzing the rs12996547 x age interaction

The rs12996547 polymorphism was not used in GTEx consortium analyses^43^. Leveraging the 1000 Genomes Project^66^ and the HaploReg database^42^, we identified a nearby SNP, rs7575617, in linkage disequilibrium (D’ = 0.89) that was used in the GTEx analyses and queried the GTEx portal for eQTL associations. Using the GTEx v8 data release, we correlated donor age (coded as a numerical variable with age in 10-year bins) and *TMEM18* gene expression within visceral adipose tissue samples.

### Screening BMI GxE interactions for pleiotropic disease associations

To test whether GxE interactions associated with BMI are also associated with related diseases, three binary disease phenotypes were assembled that represent diabetes diagnosis, high blood pressure diagnosis (HBP), and coronary artery disease ascertainment (CAD). Diabetes and HBP was coded using corresponding fields 2443-0.0 and 6150-0.0, which include self-reported questionnaire information. The diabetes phenotype represents a self-reported answer to the question, “Has a doctor ever told you that you have diabetes?” This would represent a mix of diabetes subtypes, including Type 1 and Type 2 diabetes. For diabetes, values less than 0 were removed from association testing. For HBP, value less than 0 represented controls and value equal to 4 represented a case. CAD was specified using criteria from previous research^22^. The following individuals were listed as cases: field 20002-0.0 equal to 1075; fields 41203-0.0 or 41205-0.0 equal to 410, 4109, 411, 4119, 412, 4129; fields 41202-0.0 or 41204-0.0 equal to I21, I210, I211, I212, I213, I214, I219, I21X, I22, I220, I221, I228, I229, I23, I230, I231, I232, I233, I234, I235, I236, I241, I252; fields 41200-0.0 or 41210-0.0 equal to K40, K401, K402, K403, K404, K41, K411, K412, K413, K414, K451, K452, K453, K454, K455, K491, K492, K498, K499, K502, K751, K752, K753, K754, K758, K759. All other individuals were listed as the controls for CAD.

GxE interactions with *FDR* < 0.1 and same direction of effect in discovery and replication sets were tested for association with diabetes, HBP, and CAD risk. *P*_R_ < 0.05 in the replication cohort was not required, because the limited sample size in the replication cohort (one quarter the size of the discovery set) may reduce power to identify interaction associations at that level of significance and typically implies that a larger effect needs to be observed within the smaller cohort to reach that level of significance. (We found that only requiring direction of effect will still show statistically significant differences in replication rate between QTLs and random genome-wide SNPs.) Finally, the GxE interaction with disease was tested for by employing logistic regression with identical covariates to the BMI analysis. This was repeated with adjustment for BMI by using BMI as a covariate.

### PheWAS enrichment of vQTLs using Open Targets database queries

We identified all SNP-phenotype associations with *P* < 0.05 and tested whether raw vQTLs were enriched for certain phenotype associations compared to pure muQTLs (Methods).

To determine whether variance QTLs were more likely to be associated with some phenotypes, a phenotype-wide association study (PheWAS) leveraging the Open Targets database was performed. For an input SNP, we identified all phenotypes within the database that have been associated using a previous GWAS at *P* < 0.05. Across a set of queried SNPs, we calculated the proportion that were associated with the phenotype. We repeated this for the set of variance QTLs and the set of pure mean QTLs (no vQTL association). We trimmed the phenotypes list using the Open Target categories that were relevant to our study (Supp Data 7).

Next, we used a statistical test to determine whether a given set of SNPs is enriched for association with a phenotype compared to a background set. Given a test set of *K* SNPs, in which *m* of the *K* SNPs are associated with the phenotype, and a background SNP set in which *p* is the proportion of SNPs associated with phenotype, we employ an exact binomial test with *m* successes, *K* trials, and *p* hypothetical probability of success. We test significance under a one-sided alternative hypothesis that the observed success rate, *m* / *K*, is greater than *p*. The *P*-values from the test were calculated for every phenotype present in the test set. False discovery rate correction was applied and significance assessed at *FDR* < 0.1.

We applied the described PheWAS enrichment test in two settings. First, we evaluated whether the test was robust for use with the real data by randomly sampling 21 pure muQTLs and using the remaining pure muQTLs as the background set. Next, we evaluated if some associated phenotypes are enriched in the vQTL set compared to in the pure muQTLs by using the pure muQTLs as the background set.

### Annotating QTLs with protein coding genes

All protein coding genes were downloaded from Ensembl. A QTL was queried in the Open Targets using the API, and the variant-to-gene (V2G) scores and the Ensembl Variant Effect Predictor (VEP) scores were saved for all Ensembl protein coding genes. If a queried SNP’s VEP score for a gene is greater than zero in Open Targets (e.g., the variant lies within an intron, exon, or UTR region), than the SNP is annotated with the gene with the greatest VEP score. Otherwise (for intergenic SNPs), the gene with the highest V2G score is used. If no protein coding genes have a V2G score (due to far proximity), the coding gene with the nearest transcription start site was identified using the Open Targets API. Finally, if the queried SNP was not present in Open Targets, then a SNP in LD was identified as a proxy (Supp Data 8). rs550990127, rs562044398, rs772168224, and rs753789664 are four indels (3 muQTLs, 1 RINT vQTL) that were removed from functional enrichment and PheWAS analyses due to annotation issues.

### GeneMania network creation and GO enrichment analysis

GeneMania incorporates multiple biological databases to create a gene network, identify highly-interconnected genes, and perform GO enrichment analysis. We used the browser platform with default settings, except for the addition of the “Attributes” databases. We queried the list of annotated genes for raw vQTLs, and separately the list of genes for pure muQTLs.

### Stratified LD-score regression to infer cell-type relevance

Stratified LD score regression was performed with gene expression data using the “Multi_tissue_gene_expr” flag and default settings. Summary statistics were transformed using the *munge_sumstats*.*py* script. Only non-MHC HapMap3 SNPs were kept for LD score regression analysis. Cell-type enrichment *P*-values across the 205 functional annotations were adjusted using the Benjamini-Hochberg method for false discovery rate^67^.

## End Notes

## Acknowledgements

We would like to thanks members of the Clark Lab and members of the Elemento Lab for helpful discussions surrounding this project. Support was provided for A.R.M. by the Tri-Institutional Training Program in Computational Biology and Medicine.

## Author Contributions

A.R.M., O.E., and A.G.C. conceived and designed the study. A.R.M performed the analysis. E.R.D., S.K., and C.V.V.H. aided in developing the methods. A.R.M., O.E., and A.G.C. wrote the manuscript. A.G.C. and O.E. supervised the study. All the authors reviewed and approved the manuscript.

## Declaration of Interests

O.E. is scientific advisor and equity holder in Freenome, Owkin, Volastra Therapeutics and One Three Biotech. C.V.V.H. is an employee of the Regeneron Genetics Center.

## Data and Code Availability

UK Biobank data was accessed under application number 47137. Code to run a Deviation Regression Model on any genome-wide association study data is available at https://github.com/drewmard/DRM. Computer code to reproduce the analyses are available at https://github.com/drewmard/ukb_vqtl.

