## Supplementary Information for "Leveraging phenotypic variability to identify genetic interactions in human phenotypes"

**Marderstein et al.**

### SUPPLEMENTARY FIGURES

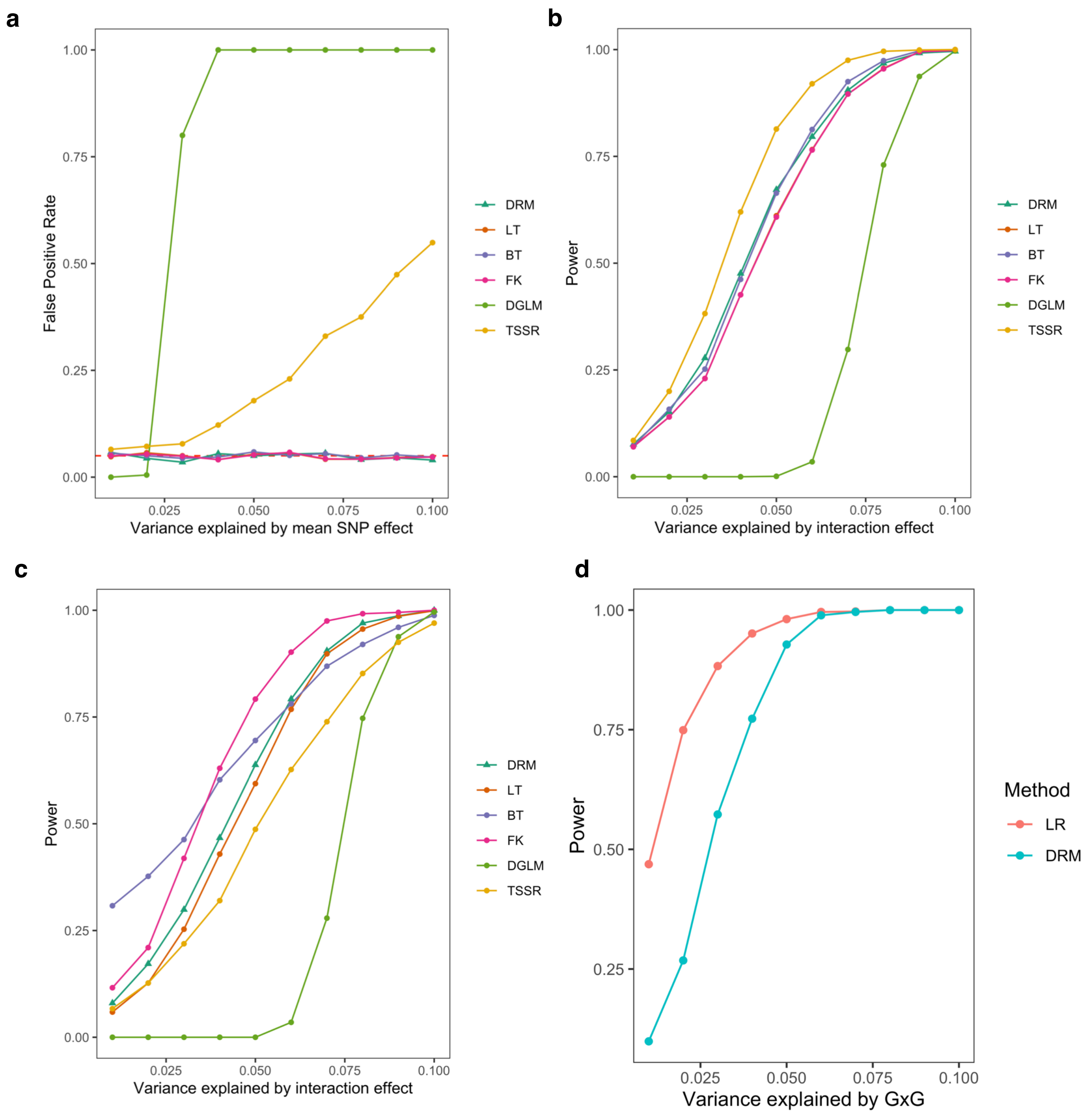


**Supplementary Figure 1: FPR and power of variance tests.** (a) FPR for different variance tests under variable mean QTL effect sizes in a normal distributed phenotype. (c-d) Power comparison between variance tests in normally distributed phenotypes (c) and chi-square phenotypes (d) as a function of varied interaction effect sizes. Deviation Regression Model (DRM), Levene's test (LT), Bartlett's test (BT), Fligner Killeen test (FK), double generalized linear models (DGLM), two-step squared residual approach (TSSR). FPR and power calculated as the proportion of simulations with $P < 0.05$. (d) Power of muQTL and vQTL tests to identify interaction SNPs. An increase in the proportion of variance explained by the interaction (x-axis) leads to an increase in power to detect interaction SNPs for both muQTL approaches (red, using linear regression (LR)) and vQTL approaches (blue, using the Deviation Regression Model (DRM)), as measured across 1000 simulations. Across all interaction effect sizes, the linear regression approach has higher power than the Deviation Regression Model. However, any muQTLs would also be detected. Therefore, any SNPs that directly influence the phenotype with no interactions will be picked up as well.

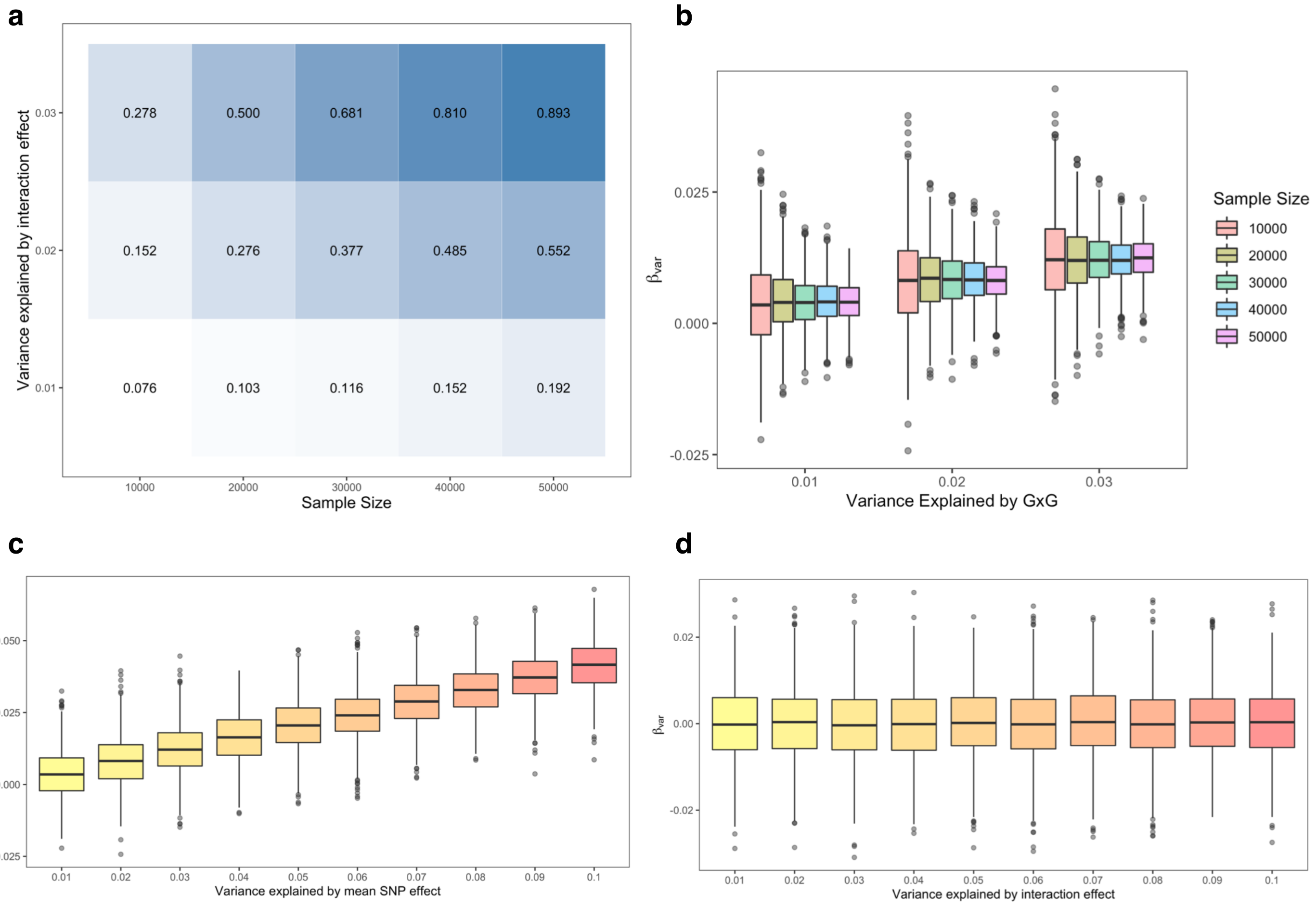


**Supplementary Figure 2: DRM power as a function of sample size and effect size.** (a) Effect size is the proportion of phenotypic variance due to the interaction (y-axis). Light to dark color reflects low to high statistical power. Power calculated across 1000 simulations. (b) Influence of sample size on estimated variance effects. The standard error of the DRM-estimated variance effect (y-axis) is decreased with larger sample sizes (boxplot grouping). Results shown for 1%, 2%, and 3% variation explained by the interaction (x-axis). (c-d) Influence of interaction effects on estimated variance effects. (c) An increase in the variation explained by an interaction (x-axis) leads to an increase in the DRM-estimated variance effect (y-axis). (d) However, an increase in the variation explained by a SNP influencing the mean but not the variance of a phenotype (x-axis) does not lead to an increase in the DRM-estimated variance effect (y-axis). In (b-d), results are summarized and displayed in boxplots. Each point represents a new simulation iteration. The middle line is the median, the lower and upper hinges represent first and third quartiles, and the whiskers extend from the hinge with a length of 1.5x the inter-quartile range. All data points outside the lower and upper whisker ends are shown individually.

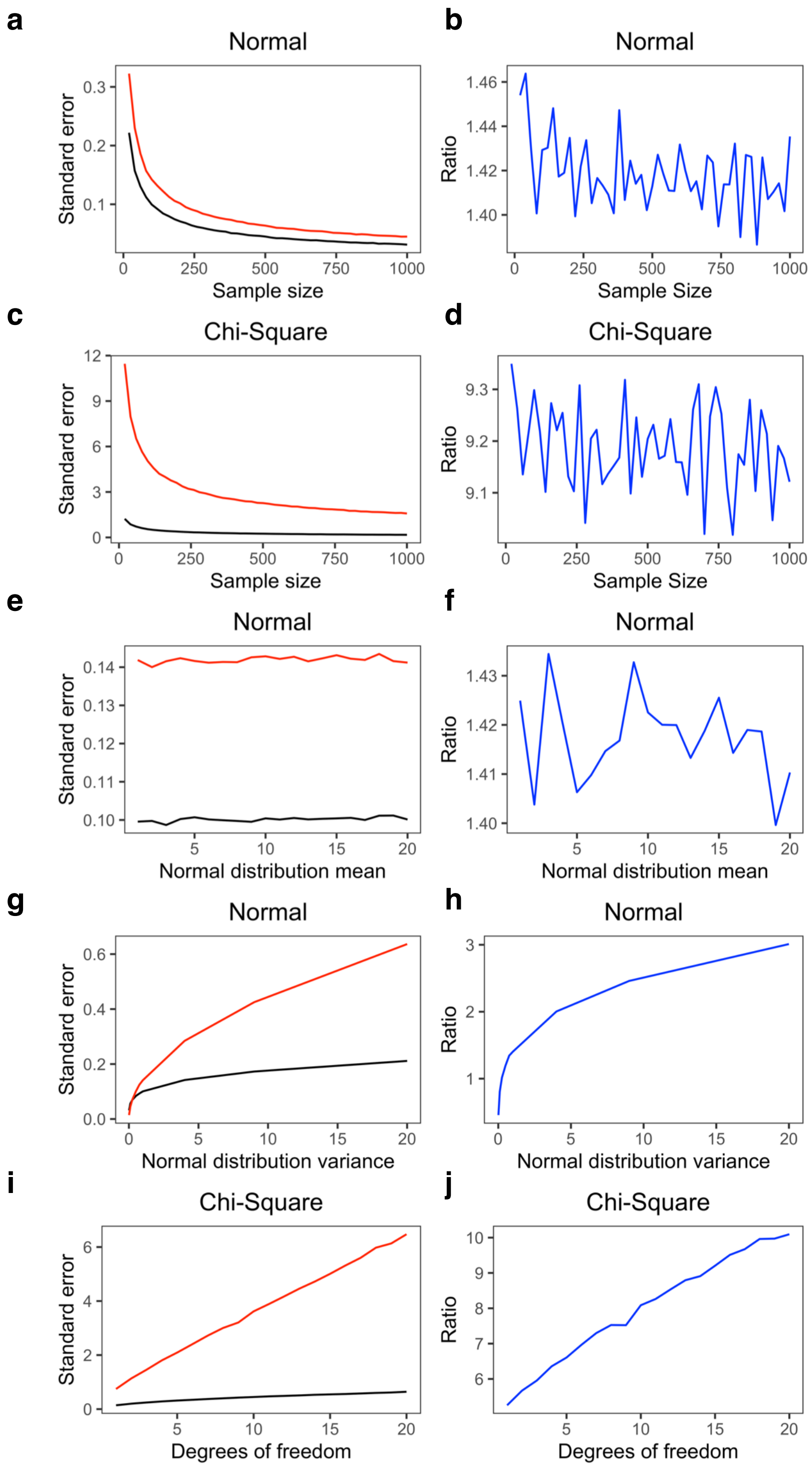


**Supplementary Figure 3: Standard error of the mean and variance.** Standard error of the mean and variance estimates in normal and chi-square distributions, as a function of either sample size (a-d) or distribution parameters (e-j). In the left column figures, the red line describes the standard error of variance ($\sigma_\sigma$) and the black line describes the standard error of mean ($\sigma_\mu$). The ratio $\sigma_\sigma$ / $\sigma_\mu$, which describes how the variance standard error changes relative to the mean standard error, is displayed in the right column figures as a blue line.

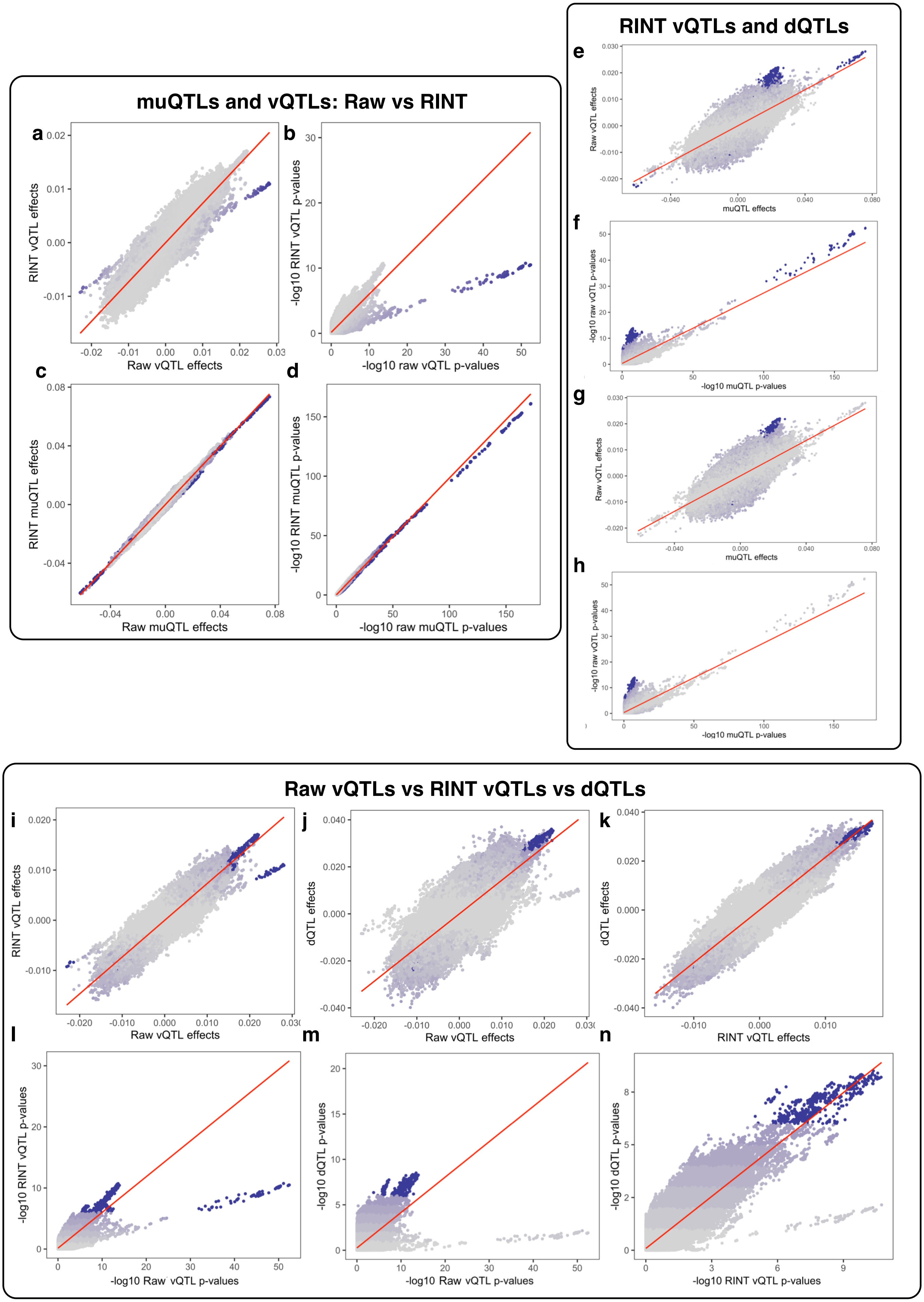

muQTLs and vQTLs: Raw vs RINT
a
b
c
d
RINT vQTL effects
Raw vQTL effects
-log10 RINT vQTL p-values
-log10 raw vQTL p-values
RINT muQTL effects
Raw muQTL effects
-log10 RINT muQTL p-values
-log10 raw muQTL p-values
RINT vQTLs and dQTLs
e
f
g
h
Raw vQTL effects
muQTL effects
-log10 raw vQTL p-values
-log10 muQTL p-values
Raw vQTLs vs RINT vQTLs vs dQTLs
i
j
k
l
m
n
RINT vQTL effects
Raw vQTL effects
dQTL effects
RINT vQTL effects
-log10 RINT vQTL p-values
-log10 Raw vQTL p-values
-log10 dQTL p-values
-log10 RINT vQTL p-values

**Supplementary Figure 4: The different ”flavors” of QTLs.** (a-d) Influence of raw and RINT analyses on muQTLs and vQTLs. A genome-wide association study was performed to search for mean (muQTL) and variance (vQTL) effects on body mass index (BMI) levels. It was repeated for both rank inverse normal transformed (RINT) and untransformed BMI (raw). (a-b) Estimated effect sizes (a) and *P*-values (b) between raw vQTL and RINT vQTL analyses. Colors represent $-\log_{10}$ raw muQTL *P*-values. (c-d) Estimated effect sizes (c) and *P*-values (d) between raw muQTL and RINT muQTL analyses. Colors represent $-\log_{10}$ raw vQTL *P*-values. Red lines represent line of best fit. (e-h) RINT-based QTLs are displaced from a mean-variance relationship. A genome-wide association study (GWAS) was performed to search for mean (muQTL) and variance (raw vQTL) effects on untransformed body mass index (BMI) levels. A GWAS was also performed on the rank inverse normal transformed (RINT) BMI to search for variance (RINT vQTL) and dispersion (dQTL) effects. Estimated effect sizes (e, g) and $-\log_{10}$ p-values (f, h) are shown for the muQTL and raw vQTL analyses, with light to dark colors representing $-\log_{10}$ RINT vQTL p-values in panels e-f and $-\log_{10}$ dQTL p-values in g-h. SNPs with $P < 10^{-5}$ colored in dark purple. The dQTLs (bottom row) are slightly more displaced from the general mean-variance relationship compared to the RINT vQTLs. These figures can be compared to those in Main Text Figures 3. (i-n) Relationship between raw vQTLs, RINT vQTLs, and dQTLs. Comparison of raw vQTLs, RINT vQTLs, and dQTLs according to effect sizes (i-k) and significance (l-n). The red line represents the line of best fit. Points are colored by the $-\log_{10}$ p-value of the y-axis analysis, with purple representing significant ($P < 5 \times 10^{-8}$ with raw BMI, $P < 10^{-5}$ with RINT BMI).

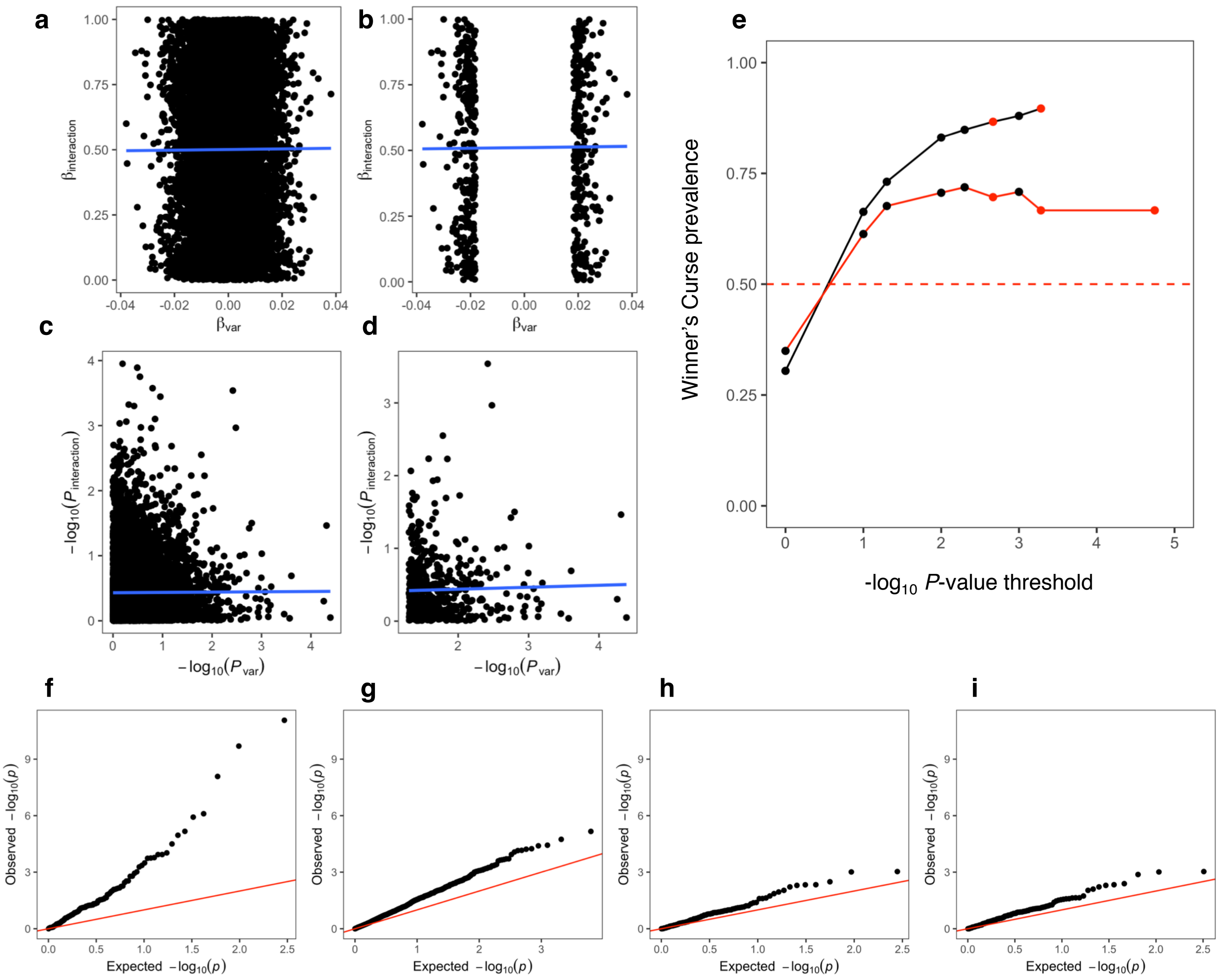


**Supplementary Figure 5: Discovering GxE interactions.** (a-d) False positive vQTLs do not correlate with false positive interactions. 10,000 simulations of a SNP-by-factor interaction were performed. For each iteration, statistical testing was performed between the SNP and the variance of the phenotype, and the SNP-factor interaction and the mean of the phenotype. The relationship between interaction and variance effect sizes (a-b) and *P*-values (c-d) are shown. In (b) and (d), the data is subset to the simulations where the variance *P*-values are below 0.05. (e) Winner's curse is reduced at QTLs. Winner's curse describes the upward bias in the magnitude of effect sizes within significant associations from a GWAS. The prevalence of winner's curse in the GxE analyses were calculated by estimating the proportion of interactions where the magnitude of the estimated effect size in the discovery cohort was larger than the magnitude of the estimated effect size in the replication cohort. This was calculated for all interactions where the direction of effect was identical. In the figure, given a threshold *x* (x-axis), the winner's curse prevalence (y-axis) is calculated for all interactions with $P_D < x$. The upper black line indicates GxE interactions using matched genome-wide SNPs. The lower red line indicates GxE interactions using all QTL-nominated SNPs. The expected winner's curse prevalence under random observations (50%) is shown as a horizontal dashed red line. Red points are $FDR < 0.1$, $< 0.05$, and $< 0.01$ cut-offs. (f-i) Discovery of GxE interactions using different QTL criteria. Quantile-quantile plots for GxE interactions across seven environmental factors using (f) 21 raw vQTLs, (g) 448 muQTLs that were not raw vQTLs, (h) 20 RINT vQTLs that were not raw vQTLs, or (i) 23 dQTLs that were not raw vQTLs. The *x*-axis shows the $-\log_{10}$ *p*-values under the null distribution and the *y*-axis shows the observed $-\log_{10}$ *p*-values, where each point represents a different GxE interaction. The red line represents the expectation under the null, with intercept = 0 and slope = 1.

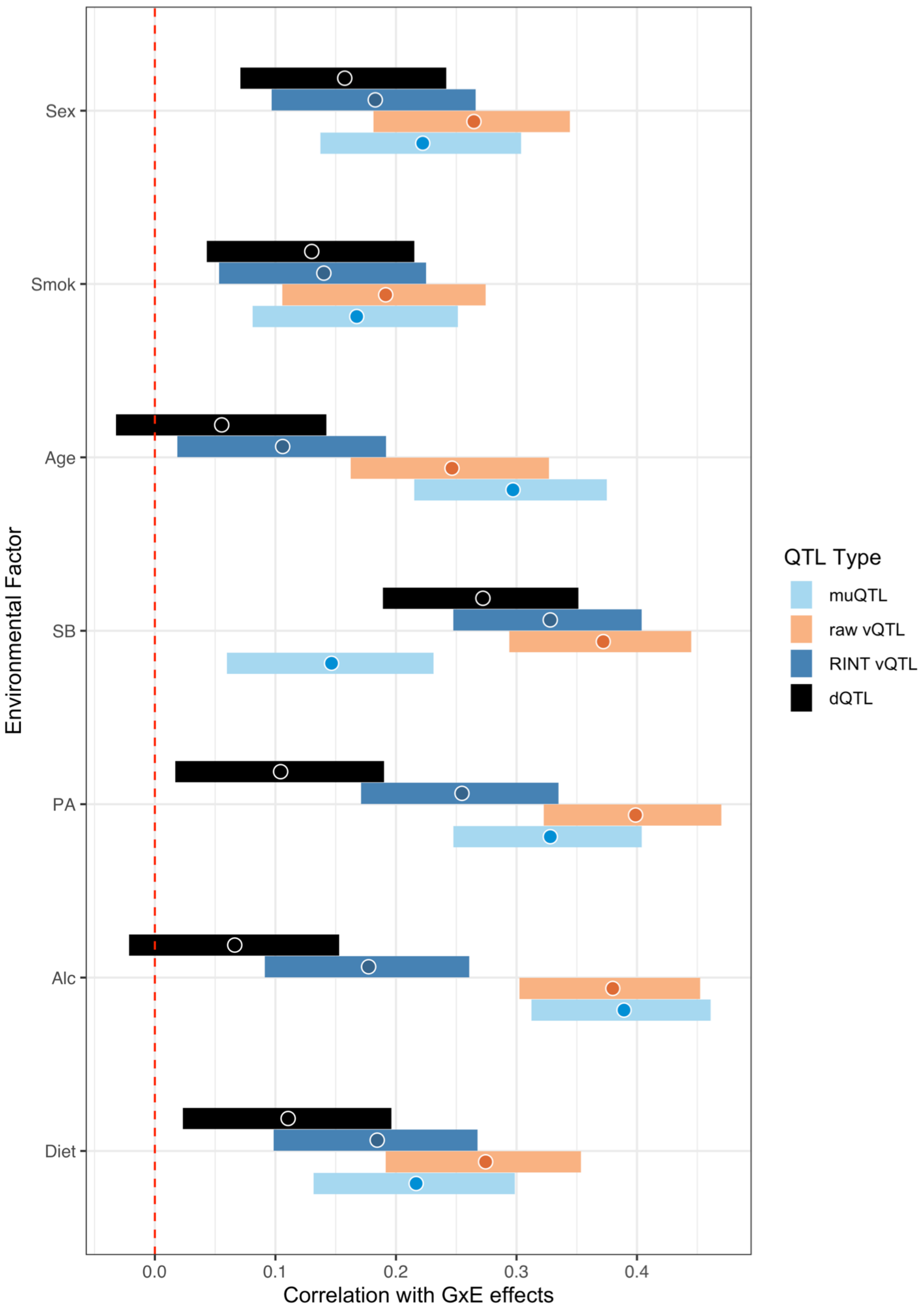


**Supplementary Figure 6: Marginal QTL effects correlate with estimated GxE effects.** The estimated marginal effects from the muQTL (light blue), raw vQTL (golden), RINT vQTL (dark blue), and dQTL (black) analyses were correlated with the estimated GxE effects for each co-factor separately. The estimated Spearman's rho is shown with 95% confidence intervals based on standard errors. Raw vQTL effects are either the 1st or 2nd best correlated QTL effect with the estimated GxE effects across all environmental factors. MuQTLs had the weakest correlation with GxE effects using sedentary behavior.

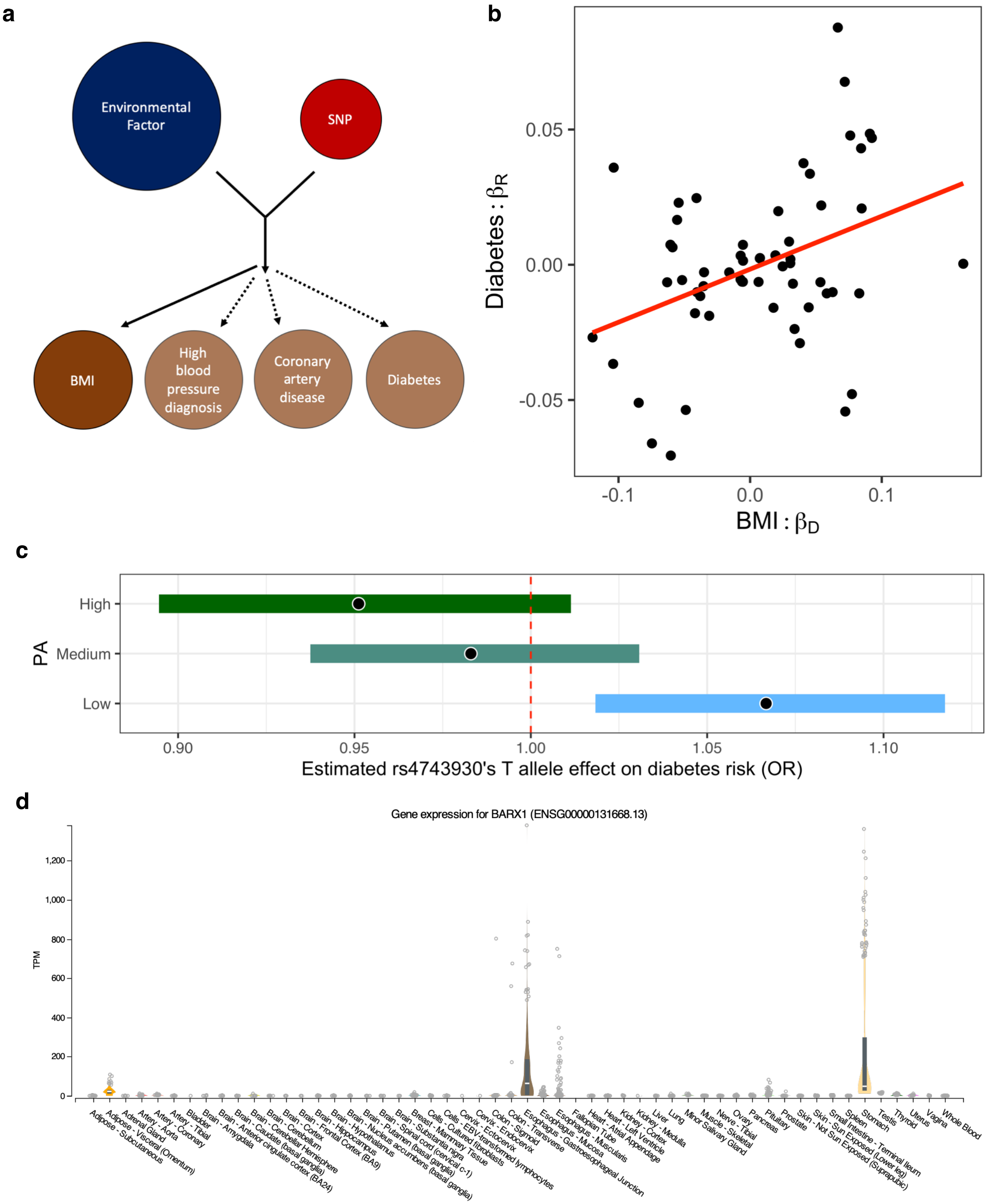


**Supplementary Figure 7: Pleiotropic GxE effects.** (a) Schematic diagram. The original analysis identified GxE interactions associated with BMI. Using a PheWAS-inspired approach, we tested for association between GxE interactions and high blood pressure diagnosis risk, coronary artery disease risk, and diabetes risk. (b) Estimated GxE effects in BMI within the 80% discovery set (x-axis) from linear regression were correlated with estimated GxE effects on diabetes risk within the 20% replication set (y-axis) from logistic regression. Each data point represents a different SNP x co-factor interaction. In contrast to Figure 5b, the estimated GxE effects on diabetes risk were adjusted for measured BMI. BMI GxE interactions appear predictive of diabetes GxE interactions, even after accounting for BMI. (c) The estimated marginal effect of the rs4743930 T allele on diabetes risk, conditioned on physical activity (PA) levels and adjusted for measured BMI. Estimated diabetes risk effect is in terms of the relative odds ratio (OR). The estimate is shown by the black dot, and the bars indicate the 95% confidence intervals. (d) *BARX1* gene expression across GTEx tissues. Figure made by the GTEx Consortium.

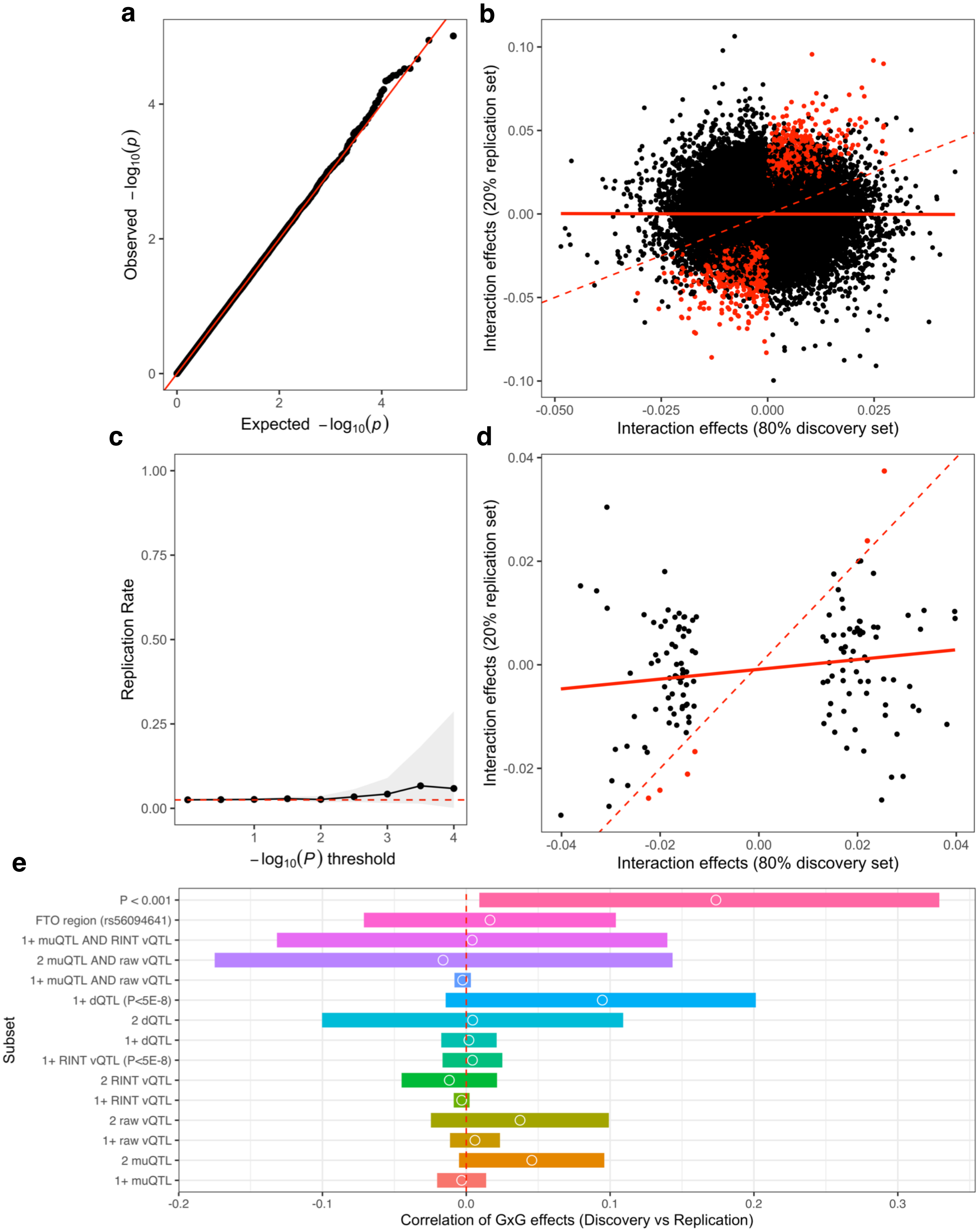


**Supplementary Figure 8: Searching for GxG interactions.** (a) Quantile-quantile plot showing the observed $-\log_{10}(P)$ for all-pairwise interactions based on QTLs (y-axis) compared to the theoretical $-\log_{10}(P)$ distribution (x-axis). The red line has slope 1 and intercept 0, indicating the expected p-values under the null distribution. There does not appear to be any deviation from the expectation. (b) Relationship between estimated GxG effects within the discovery cohort and the replication cohort. Each data point represents a different GxG interaction, and are colored red if replicated within the replication cohort (according to same direction of effect and $P_R < 0.05$). (c) GxG interactions, as quantified by those with the same direction of effect in both discovery and replication sets and $P_R < 0.05$. Given a threshold $x$ (x-axis), the replication rate (y-axis) is calculated for all interactions with $P_D < x$. The grey area represents the confidence interval from a binomial test with hypothesized replication rate equal to 0.025. (d) Same figure as (b) except interactions are subset to those with $P_{80} <$ 0.001. (e) The estimated GxG effect sizes within the discovery set was correlated with the estimated GxG effect sizes within the replication sets (y-axis). The set of GxG interactions was subset according to the label along the “Subset” axis, where “1+” indicates that the GxG interactions contain at least 1 of that QTL type, “2” indicates both SNPs in the interaction are from that QTL type, “P<5E-8” indicates a stricter significance threshold of $P < 5 \times 10^{-8}$, “AND” indicates the SNP must pass both QTL type criteria. “FTO region” indicates the 501 interactions between rs56094641 (the *FTO* intron region’s tag SNP in our study) and the 501 other identified QTLs. “P < 0.001” specifies the GxG interactions with $P < 0.001$ in the discovery cohort. The estimated correlation is shown by the white outlined circle, while the confidence interval of this estimate is outlined by the minimum and maximum values of the bar. Correlation equal to 0 is outlined by a red line.

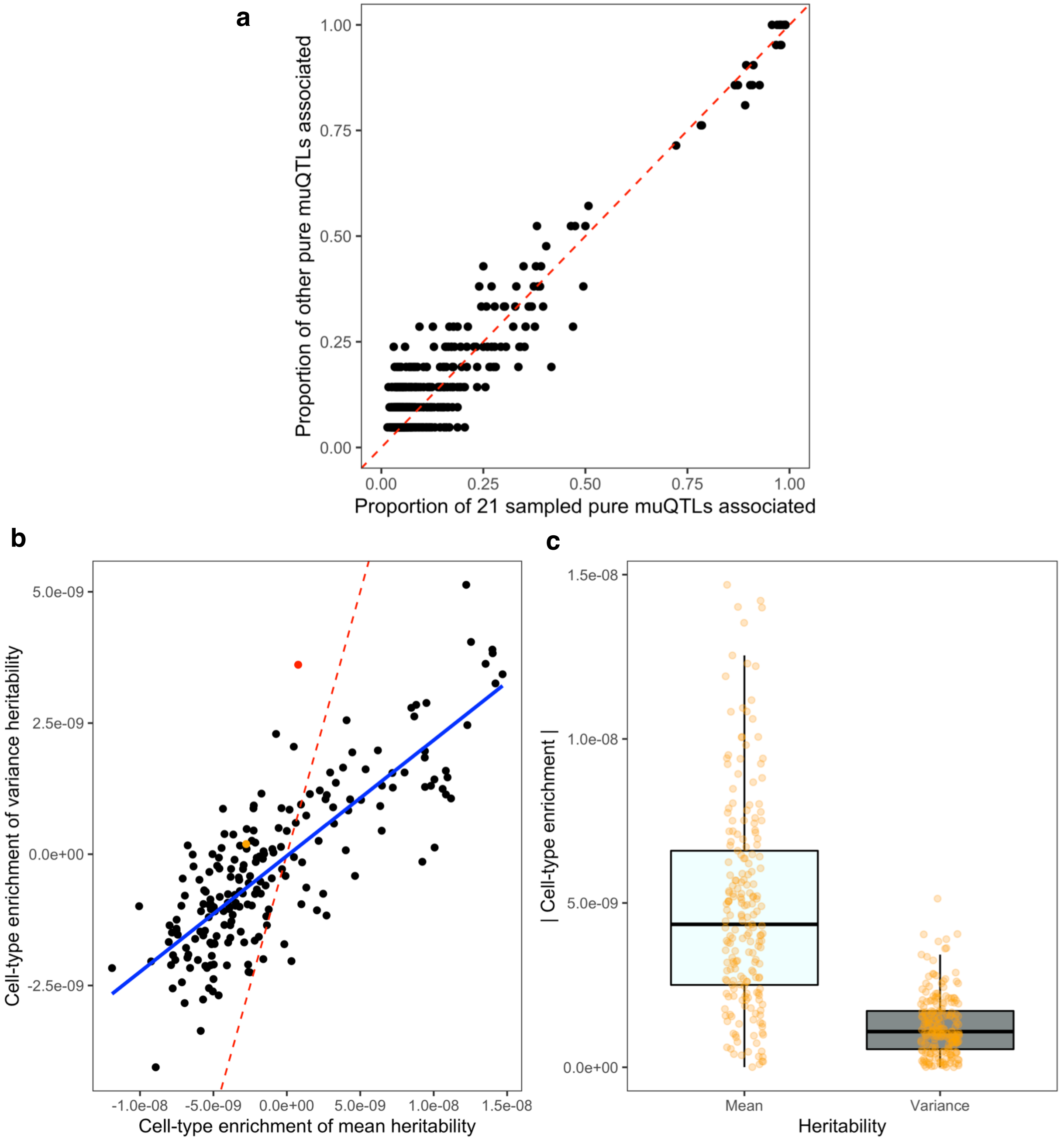


**Supplementary Figure 9: PheWAS enrichment and partitioned LD-score regression**. (a) Control experiment for the PheWAS enrichment test. A PheWAS was performed using the Open Targets database with P < 0.05 and a binomial test is used to determine whether a phenotype is more often associated within a test set of SNPs compared to a background set of SNPs. The proportion of SNPs in the test set that are associated with a given phenotype is shown along the y-axis, and proportion of SNPs in the background set that are associated with a given phenotype is shown along the x-axis. Each point represents a different phenotype in the Open Targets database, where a red colored point indicates *FDR* < 0.1. In this figure, the test set is a random set of 21 pure muQTLs (muQTLs with no vQTL significance), and the background set is the remaining set of pure muQTLs. (b-c) Stratified linkage disequilibrium score regression was performed on the muQTL summary statistics and the raw vQTL summary statistics using the "Multi_tissue_gene_expr" gene set. (b) The enrichment coefficients for the mean (x-axis) and variances (y-axis). Each data point represents a different cell-type category. The points colored red and orange represent the "A03.556.875.875.Stomach" (Franke data) and "Stomach" (GTEx data) categories respectively. The red line has slope equal to 1 and intercept equal to 0, as if mean and variance heritability enrichments were identical. The blue line is the line of best fit. c) The magnitude of cell-type enrichments in mean and variance analyses. Magnitude is the absolute value of the cell-type category coefficient. Each point represents a different cell-type category. In the box plots, the middle line is the median, the lower and upper hinges represent first and third quartiles, and the whiskers extend from the hinge with a length of 1.5x the inter-quartile range. The magnitude of cell-type enrichments in mean BMI heritability is larger than the enrichments in variance heritability ($P = 3.6 \times 10^{-45}$). This is expected as the mean GWAS signal was much stronger than the variance signal.

### SUPPLEMENTARY NOTES

#### Supplementary Note:

##### Supplementary Note 1: Influence of sample size and interaction effects on $\beta_{var}$

In our simulations for comparing variance test methods, we also varied the sample sizes and interaction effects and assessed the influence on estimated variance effects ($\beta_{var}$) and power from the DRM. As shown in Supplementary Figure 2, the power increases as either sample size or interaction effects increase. For example, the power to detect a $V_G$ = 2% interaction in an *N* = 10,000 population using the DRM is similar to the power to detect a $V_G$ = 1% interaction in an *N* = 40,000 population (Supplementary Figure 2a).

The improved DRM power from an increase in sample size is due to the standard error of the variance effects $\beta_{var}$ reducing, as $\beta_{var}$ becomes more consistent with less variability (Supplementary Figure 2b). Due to an increase in interaction effect sizes, $\beta_{var}$ increases proportionally to the size of interaction effects (Supplementary Figure 2c). This is contrast to the FPR simulations, where there was no correlation between single-SNP mean effects and $\beta_{var}$ at SNPs with only a pure mean effect (Supplementary Figure 2d).

##### Supplementary Note 2: Mean GWAS comparison to Neale Lab analysis

We confirmed our mean-based GWAS results by observing that the -log10 *P*-values recapitulated the -log10 *P*-values from the publicly available Neale Lab analysis[1] ($r$ = 0.93). However, we had decreased power, with the most significant associations in the Neale Lab analysis having lower *P*-values than in our analysis. We attributed this to reduced sample size in our analysis from having a held-out validation set containing 68,840 individuals.

##### Supplementary Note 3: Mean and variances are correlated in non-normal distributions

In a sample from a normal distribution, the variance, an ancillary statistic, is independent from the mean, a sufficient statistic, via Basu's theorem[2]. The normal distribution is the only statistical distribution where the sample mean and sample variance are independent. No other distribution has this property.

In a non-normal distribution, the distributional parameters jointly influence both the mean and variance. For example, the chi-square distribution is characterized by a single parameter *k*, also known as the degrees of freedom. The parameter *k* determines the number of squared normal random variables that are jointly summed to create the chi-square distribution. The mean and variance of the chi-square distribution is equal to *k* and *2k* respectively. Therefore, any change in *k* will also impact a sample's mean and variance. It can be implied that any genetic effect which influences the means of a non-normal distribution will also have an effect on the variances through this general relationship.

##### Supplementary Note 4: Standard error of variance estimates versus mean estimates

We identified many fewer vQTLs compared to muQTLs in our analyses. Using simulation, show that this is due to a typically much higher standard error in sample variance compared to the standard error in sample means.

We first show that the standard errors of mean and variance estimates are greatly influenced by sample size. We generated *N* data points from a standard normal distribution (mean = 0 and variance = 1) and calculated the mean and variance of the *N* sampled values. We repeated this data generation process 10,000 times for each value of *N*, and estimated the standard error of the mean and variance estimates by calculating the standard deviation of the sample estimates across all 10,000 sets. We iterated over all increments of 20 for $N = 20$ to $N = 1{,}000$ data points, calculating the standard deviation of the mean estimates ($\sigma_{\mu}$) and the standard deviation of the variance estimates ($\sigma_{\sigma}$) at each $N$ value. We found that both $\sigma_{\mu}$ and $\sigma_{\sigma}$ decrease as sample size decreases (Supplementary Figure 3a). However, the ratio $\sigma_{\sigma} / \sigma_{\mu}$, which describes how the variance standard error changes relative to the mean standard error, stayed relatively similar across values of $N$ (Supplementary Figure 3b). We had the same observations when using $N$ data points from a chi-square distribution with 15 degrees of freedom (Supplementary Figures 3c-d). This data suggests that an increase in sample size leads to improved accuracy for estimating means and variances, but an increase in sample size (given the distributional parameter settings) will not lead to having superior accuracy for the estimation of variances compared to the estimation of means.

We next analyzed how the parameters of the underlying distribution influence $\sigma_{\mu}$ and $\sigma_{\sigma}$. First, we repeated the process above for a normal distribution, except keeping $N = 100$ and iterating over mean = 1 to 20 in 1 unit increments (keeping the variance constant at 1). We found that the true mean value does not have a strong influence over standard error of mean or variance (Supplementary Figures 3e-f). We then generated data by keeping constant $N = 100$ and constant mean = 0, but at different variance values: 0.01, 0.1, 0.25, 0.5, 0.75, 1, 4, 9, and 20. We found that an increase in the true variance would increase $\sigma_{\sigma}$ and $\sigma_{\mu}$, although with an interesting relationship. When the true variance was very small (reduced to below 0.25), the estimated standard error for variance was smaller than the estimated standard error for mean ($\sigma_{\sigma} / \sigma_{\mu} < 1$ ). $\sigma_{\sigma}$ increased with a greater rate than $\sigma_{\mu}$, and thus $\sigma_{\sigma} / \sigma_{\mu} > 1$ for larger true variance values (> 0.5) (Supplementary Figures 3g-h). This suggests that if the underlying data has very low variability, variance estimates may be more accurate to estimate than mean estimates. As the previous results suggested and we further confirm, this relationship is independent of sample size. However, in more typical data distributions with larger variability, variance estimates are more difficult to accurately estimate.

Finally, we further show that in a chi-squared distribution (where the degrees of freedom specify the mean and the variance of the non-normal distribution) with degrees of freedom selected at the integers between 1 and 20, $\sigma_{\sigma} / \sigma_{\mu} > 1$ and continues to increase as the degrees of freedom increases (Supplementary Figures 3i-j).

Overall, we show that variance estimates are more difficult to accurately estimate (have higher standard error) than mean estimates (which have lower standard error), unless there is very little variability in the distribution. Therefore, in the context of genetic studies, attempting to detect differences in phenotypic variance estimates between genotypes will be far more difficult compared to the identification of differences in phenotypic mean estimates between genotypes.

**Supplementary Note 5: Statistical methods and transformations under a mean-variance relationship**

Young et al.[3] previously described how a rank inverse normal transformation (RINT) decorrelates mean and variance effects and a subsequent dispersion effect test (DET) can infer robust changes in phenotypic variance independent of genotypic changes and any phenotypic transformations. They implemented a heteroskedastic linear mixed model (HLMM), which jointly estimates mean and log-linear variance effects, to perform the DET. Using these statistical tools, we performed a comprehensive analysis of mean, variance, and dispersion effects in untransformed and RINT body mass index levels, contrasting the Young et al. approaches with our own. We refer to mean (linear regression, or LR) and variance (DRM) effects on the untransformed phenotype as “Raw muQTLs” and “Raw vQTLs”; mean (LR), variance (DRM), and log-linear (HLMM) effects on the transformed phenotype as “RINT muQTLs”, “RINT vQTLs”, and “RINT log-vQTLs”, and dispersion effects as “dQTLs” (DET). The dispersion effects assume that the correlation between mean and variance effects is minimal after a rank inverse normal transformation, but that a general relationship still exists.

Despite only 4 of 24 RINT vQTLs ($P < 10^{-5}$) also being genome-wide significant raw vQTLs (16.7%) ($P < 5 \times 10^{-8}$) (Figure 3e), we observed strong correlation between RINT and raw vQTL estimated effects ($r = 0.89$) (Supplementary Figure 4a). The variance *P*-values also correlated ($r = 0.68$), although it is clear that some of the RINT vQTL p-values were more conservative than the raw vQTL p-values (Supplementary Figure 4b). For example, there were only 3 independent significant RINT vQTLs compared to 21 independent significant raw vQTLs with $P < 10^{-8}$. In comparison, the mean effects were near-identical for raw and RINT BMI phenotypes (r=0.99) (Supplementary Figure 4c-d). Furthermore, we note that the RINT vQTLs and RINT log-vQTLs were very similar across the spectrum of strong and weak effects ($r = 0.94$ and $r = 0.80$ for effect sizes and p-values respectively). Therefore, we did not consider RINT muQTLs and RINT log-vQTLs in other comparisons, only using the raw muQTLs and RINT vQTLs to represent these analyses.

In our analysis, we observed a decrease in the correlation between raw muQTL and variance effects as estimated on the RINT phenotype ($r = 0.28$) compared to the untransformed phenotype ($r = 0.65$) (Figure 3b, 3d). This mean-variance relationship could be reduced further by considering the dispersion effects ($r = 0.04$) (Figure 3f), which are very similar to the RINT vQTL effects ($r = 0.91$) (Supplementary Figure 4k) but in a manner that reduces the correlation with mean effects (Figure 3f). As seen in Supplementary Figure 4e-h, the significant SNPs in either the RINT vQTL or dQTL analyses are the ones that are furthest displaced from the general mean-variance relationship in untransformed BMI (red line), with the dQTLs being more displaced from the general mean-variance correlation than the RINT vQTLs. We observed that the DET method to account for any mean effects in variance estimates resulted in only minor changes to RINT vQTL analysis, with large similarities between p-values (Supplementary Figure 4n). However, *P*-values were even more conservative in the DET framework than the RINT vQTLs (which are more conservative than the raw vQTL analysis) to subtract any mean effects. Interestingly, under the relaxed SNP-trait significance threshold of $P < 10^{-5}$ employed by the GWAS Catalog database, we found that only 4 of 27 RINT vQTLs and 1 of 26 dQTLs were also genome-wide significant ($P < 5 \times 10^{-8}$) mean QTLs (Figure 3h). As well, roughly half of the RINT vQTLs and dQTLs overlap (13 of 27 RINT vQTLs and 13 of 26 dQTLs), but there are several SNPs only identified in one analysis or the other.

Overall, by analyzing the RINT phenotype, we found loci that are not associated with phenotypic means but have some of the most significant variance effects, in contrast to analysis on the untransformed phenotype where muQTLs and vQTLs frequently overlap (Figure 3b-c). This allows identification of SNPs associated with variance distinct from the full set of raw vQTL and mean QTLs. Since RINT vQTLs and dQTLs had more conservative p-values compared to the untransformed vQTL analysis, we used a relaxed significance threshold ($P < 10^{-5}$) for the RINT vQTLs to enable identification of a greater number of loci. We include these RINT vQTLs and dQTLs in our downstream analyses as potential candidates for an interaction.

**Supplementary Note 6: RINT vQTLs reflect SNPs associated with the rank phenotype**

To analyze how a RINT decorrelates the mean and variance effects, we simulated 200,000 data points, consisting of 100,000 values from a chi-square distribution with df = 4 and 100,000 values from a chi-square distribution with df = 8 that were combined into a single dataset. The first distribution has variance = 4, and the second distribution has variance = 8.

Next, we applied a rank inverse normal transformation to the $N$ = 200,000 samples and separately analyzed the data from the df=4 distribution and the data from the df=8 distribution. We found that the data from the df=4 distribution had variance = 0.77, and the data from the df=8 distribution had variance = 0.67. This conflicts with the variance of the untransformed data, where the df=8 data has double the variance of the df=4 data.

To better understand, we transformed the data into ranks and did not include an inverse normal transformation. In the rank transformation, we found the new data to have variance = 2.5 x $10^9$ from the df = 4 data and variance = 2.2 x $10^9$ from the df = 8 data, which matches the observations from the RINT data. This data suggests that an analysis of vQTLs on ranked data is distinct from an analysis of vQTLs on untransformed data due to ranked vQTLs representing SNPs associated with the variance in rank, which can paint a very different picture from the original distribution.

**Supplementary Note 7: Differences in MAF and genotype missingness between QTLs.**

We aimed to determine whether minor allele frequency (MAF) and missing genotype data are factors that may artificially increase variance. It is possible that a lack of genotyped individuals which are minor homozygotes may lead to an increase in variance in the minor homozygote bin, and this could be one factor that leads to detecting variance QTLs.

We calculated three population-level statistics for each SNP associated through the QTL analyses: MAF, the number of individuals with non-missing genotype data, and the number of individuals that are minor homozygotes. We then performed a t-test between the different sets of QTLs (for example, compared the MAF between the set of muQTLs and the set of vQTLs). Overall, we found no difference in the mean for any population-level statistic between muQTLs, raw vQTLs, RINT vQTLs, or dQTLs ($P > 0.05$ between all set-set combinations).

In our analyses, we used these population-level statistics to match 10 genome-wide SNPs to each QTL. Given that there is no significant difference between QTL types in these attributes, we use the matched SNPs from all QTLs in analyses. By having a larger number of matched SNPs, we have greater precision in null distribution estimates.

**Supplementary Note 8: False positive vQTLs do not create false positive interactions**

Using simulations, we show how a non-causal SNP associated (by chance) with the variance of a phenotype does not have inflated false positive rates when testing for interactions (compared to SNPs not associated with variance).

We generated 10,000 simulations of 10,000 individuals, consisting of two SNPs with MAF = 0.3 (using a binomial distribution) and a normally distributed phenotype (with mean = 0 and variance = 1). We used the DRM to test for association between one of the SNPs and the variance of the phenotype, and a linear regression with a SNP-SNP interaction term to test for association between a pairwise-SNP interaction and the phenotype. There was no true causal effect of either SNP on the phenotype; the genotypes and phenotype were simulated independently.

We were interested in whether a SNP associated with the variance of the phenotype (false positive) had an increased probability for having a significant GxG interaction. Overall, we found a false positive rate = 0.053 across 526 simulations where a variance association was detected (DRM-based $P < 0.05$), which was not significantly different from the expected false positive rate = 0.05 (binomial test $P$-value = 0.69). Furthermore, the variance and GxG effect sizes and $P$-values did not correlate across the simulations ($r$ = 0.004, $P$ = 0.70 for effects; $r$ = 0.005, $P$ = 0.63 for $P$-values) or when limited to the subset of simulations with a false positive vQTL ($r = 0.009$, $P = 0.82$ for effects; $r = 0.027$, $P = 0.532$ for *P*-values) (Supplementary Figure 5a-d). In conclusion, our simulations did not suggest that the GxG false positive rate increases for false positive vQTLs.

**Supplementary Note 9: Prevalence of winner's curse.**

When only the same direction of effect was required, the validation rate was also significantly higher in the QTL set compared to the genome-wide SNP set (73.7% versus 63.8%; $P = 0.045$). Using the discovery and validation cohorts, we assessed the prevalence of "winner's curse". Winner's curse describes an upward bias in the estimated magnitude of significant SNP effects in a genome-wide association study. To calculate the prevalence in our own results, we estimated the proportion of GxE interactions where $|\beta_D| > |\beta_V|$, given the direction of effect was replicated [ $\mathrm{sign}(\beta_D) = \mathrm{sign}(\beta_V)$ ]. For the GxE interactions from the 502 unique QTLs, we found the winner's curse prevalence to be consistently steady, between 66% and 72%, as the GxE significance threshold became increasingly significant past $P_D < 0.05$. In comparison, when assessing the matched SNPs, we found that the winner's curse prevalence progressively increased from 73% to 90% as the threshold ranged from $P_D = 0.05$ to $P_D < 0.001$ (Supplementary Figure 5e). To statistically test for a difference between the prevalence of winner's curse in QTLs versus genome-wide SNPs, we used a two-sided exact binomial test to find significance $P = 5.1 \times 10^{-5}$ at $FDR_D < 0.1$ interactions (where the theoretical success rate was equal to observed winner's curse prevalence in the matched SNPs and the success and number of trials were from the QTL-derived interactions).

This indicates there is less bias of GxE estimates within the QTL results compared to genome-wide SNPs results. This could be due to a lower false positive rate for QTL-derived GxE or less reliability on needing to detect the GxE effect in the right context such that statistical significance can be achieved.

**Supplementary Note 10: GxE vQTL validation using a more stringent set of muQTLs:**

When considering a more stringent set of the 28 most significant pure muQTLs ($P_{mean} < 6.2 \times 10^{-17}$, $P_{var} > 5 \times 10^{-8}$) with a similar median mean-based *p*-value to the set of vQTLs (median $P_{mean} = 1.0 \times 10^{-21}$ in both the vQTL set and this stringent pure muQTL set), we observed a 2.7-fold enrichment for the vQTL-derived GxE validation rate (38.1% in the vQTL set versus 14.3% in the stringent pure muQTL set; $P = 0.006$). Therefore, the higher validation rate for GxE interactions at vQTLs do not appear to be due to more significant marginal mean-based *P*-values.

**Supplementary Note 11: Searching for GxG effects by leveraging QTL effects.**

As discussed in the main text, there was no correlation between interaction effects estimated in the 80% discovery set and the interaction effects estimated in the 20% validation set (r = -0.003, $P = 0.30$) (Supplementary Figure 8b). To further investigate this finding, we subset the GxG interactions and examined this correlation using 14 different criteria: (1) at least 1 muQTL involved, (2) at least 1 raw vQTL involved, (3) at least 1 RINT vQTL involved, (4) at least 1 dQTL involved, (5) both SNPs are muQTLs, (6) both SNPs are raw vQTLs, (7) both SNPs are RINT vQTLs, (8) both SNPs are dQTLs, (9) at least 1 RINT vQTL with $P < 5 \times 10^{-8}$ significance, (10) at least 1 dQTL with $P < 5 \times 10^{-8}$ significance, (11) at least 1 SNP that is both a muQTL and a raw vQTL, (12) both SNPs are muQTLs and raw vQTLs, (13) at least 1 SNP that is both a raw vQTL and a RINT vQTL, or (14) at least 1 SNP is the rs56094641 polymorphism in the *FTO* region. In all criteria settings, we found no significant correlation between estimated effects in the discovery and validation cohorts (95% confidence interval of the Spearman's rho includes 0 and $P > 0.05$) (see Supplementary Figure 8e). Therefore, leveraging QTL evidence did not provide an inroad to discovering epistatic interactions influencing BMI.

**Supplementary Note 12: Omitting the MHC region in GeneMania.**

The MHC region is quite complex, with high gene density, large linkage disequilibrium, extensive inter-individual variation, and codominance. As a result, it is difficult to map genetic loci in the MHC to genes. We decided to remove the genes from the MHC region for the purposes of annotated-gene-based functional analyses in GeneMania[4].

**Supplementary Note 13: Cell-type heritability enrichments.**

Using stratified linkage disequilibrium score regression[5], we found that the cell-type heritability enrichment for the "A03.556.875.875.Stomach" category was statistically significant in the BMI variance analysis but not significant in the BMI mean analysis. In Supplementary Figure 9b, we highlight this finding in red.

The "A03.556.875.875.Stomach" annotation is derived from data by the Franke lab. Our partitioned heritability analysis using the "Multi_tissue_gene_expr" flag also considered publicly available GTEx data[6]. We looked at the "Stomach" category, which is based on GTEx data, and found no statistical signal. However, we note that the enrichment was higher in the variance analysis compared to the mean analysis, with the enrichment positive for variances but negative for the means. (In Supplementary Figure 9b, we highlight the GTEx-based "Stomach" cell-type enrichment in orange.) Furthermore, we note that no GTEx annotations were significant in the variance analysis. All the different annotations shown in Main Text Figure 5f are derived from the Franke Lab.

### SUPPLEMENTARY TABLES

**Supplementary Table 1: *FTO* interaction effects are independent.** A model was fit with pairwise interaction terms between rs56094641, an intronic *FTO* variant, and co-factors smoking status (Smoking), age, diet, sedentary behavior (SB), physical activity level (PA), and alcohol intake frequency (Alc). The effect size, standard error, test statistic, and p-value of the interaction term is shown in columns 2-5.

| Interaction Term | Estimate | Std. Error | t value | Pr(>\|t\|) |
|---|---|---|---|---|
| SNP*Smoking | 0.06222388 | 0.02867939 | 2.16963751 | 0.03003545 |
| SNP*Age | -0.0059631 | 0.00176839 | -3.3720501 | 0.00074625 |
| SNP*Diet | 0.04077586 | 0.01460421 | 2.79206302 | 0.0052378 |
| SNP*SB | 0.02001741 | 0.00580839 | 3.44629093 | 0.00056845 |
| SNP*PA | -0.0796201 | 0.0181919 | -4.3766805 | 1.21E-05 |
| SNP*Alc | 0.04830689 | 0.00956766 | 5.04897698 | 4.45E-07 |

### SUPPLEMENTARY DATA

**Legends**

**Supplementary Data 1: Ensembl Gene IDs for raw vQTLs.** SNPs associated with the variance of untransformed BMI. Genes were annotated using Open Targets[7] (see Methods). SNPs in the MHC and their corresponding genes have been removed.

**Supplementary Data 2: Ensembl Gene IDs for pure muQTLs.** SNPs associated with the mean but not the variance of untransformed BMI. Genes were annotated using Open Targets (see Methods). SNPs in the MHC and their corresponding genes have been removed.

**Supplementary Data 3: GeneMania report for raw vQTL genes.** List of genes were input into GeneMania. The output data includes information on the network nodes (genes) and edges (gene networks used).

**Supplementary Data 4: GeneMania-based GO enrichment for raw vQTL genes.** List of gene ontology (GO) terms that were significant within the GeneMania network. Listed are all GO terms with $FDR < 0.1$. “Genes in network” represent the number of genes in both the GeneMania network and the GO term’s gene list, while “Genes in genome” is the gene count within the GO term’s gene list.

**Supplementary Data 5: GeneMania report for pure muQTL genes.** List of genes were input into GeneMania. The output data includes information on the network nodes (genes) and edges (gene networks used).

**Supplementary Data 6: GeneMania-based GO enrichment for pure muQTL genes.** List of gene ontology (GO) terms that were significant within the GeneMania network. Listed are all GO terms with $FDR < 0.1$. “Genes in network” represent the number of genes in both the GeneMania network and the GO term’s gene list, while “Genes in genome” is the gene count within the GO term’s gene list.

**Supplementary Data 7: Kept phenotype categories in Open Targets.** List of phenotype categories that were kept for downstream PheWAS enrichment analyses. These phenotype categories were curated by Open Targets, and each phenotype in Open Targets is assigned to a single phenotype category.

**Supplementary Data 8: Proxy SNPs in LD as replacement SNPs.** Some SNPs nominated as the lead SNP within our analyses were not present in the Open Targets database. These missing SNPs (column 1) were re-assigned to proxy SNPs in linkage disequilibrium (column 2). Four SNPs were not able to be re-assigned due to annotation issues; these are marked.

**Supplementary Data 9: Full PheWAS enrichment table.** All phenotypes that are significantly enriched ($FDR < 0.1$) in the raw vQTLs compared to the pure muQTLs. muQTL and vQTL columns specify the proportion of SNPs associated with the phenotype in each respective set. Diff column is the difference in proportions, and vQTL/muQTL is the ratio between proportions. PVAL is the nominal *P*-value and FDR is the multiple test-corrected significance value (via Benjamini-Hochberg false discovery rate[8]).

**Supplementary Data 10: QTL summary statistics.** The 502 unique QTLs from the BMI GWAS. Columns, left to right: SNP ID, chromosome, position, reference allele, minor allele, minor allele frequency, effect size and P-value of mean GWAS on untransformed BMI (linear regression), effect size and P-value of variance GWAS on untransformed BMI (DRM), effect size and P-value of variance GWAS on transformed BMI (DRM), and effect sizes and P-values of Young et al.'s GWAS approach on transformed BMI (HLMM-derived additive, log-linear variance, additive-variance, and dispersion statistics and P-values).

#### Access

Supplementary Data files can be accessed at the following link:
https://drive.google.com/drive/folders/1aarsyReLNuQjYEEMzmwnI-gNygLKFApz?usp=sharing
